# Histone variant H2BE enhances chromatin accessibility in neurons to promote synaptic gene expression and long-term memory

**DOI:** 10.1101/2024.01.29.575103

**Authors:** Emily R. Feierman, Sean Louzon, Nicholas A. Prescott, Tracy Biaco, Qingzeng Gao, Qi Qiu, Kyuhyun Choi, Katherine C. Palozola, Anna J. Voss, Shreya D. Mehta, Camille N. Quaye, Katherine T. Lynch, Marc V. Fuccillo, Hao Wu, Yael David, Erica Korb

**Affiliations:** Neuroscience Graduate Group, University of Pennsylvania Perelman School of Medicine, Philadelphia, PA; Cell and Molecular Biology Graduate Group, University of Pennsylvania Perelman School of Medicine, Philadelphia, PA; Department of Genetics, University of Pennsylvania Perelman School of Medicine, Philadelphia, PA; Epigenetics Institute, University of Pennsylvania Perelman School of Medicine, Philadelphia, PA; Department of Neuroscience, University of Pennsylvania Perelman School of Medicine, Philadelphia, PA; Chemical Biology Program, Memorial Sloan Kettering Cancer Center; Tri-institutional PhD Program in Chemical Biology, New York, NY

## Abstract

Regulation of histone proteins affects gene expression through multiple mechanisms including exchange with histone variants. However, widely expressed variants of H2B remain elusive. Recent findings link histone variants to neurological disorders, yet few are well studied in the brain. We applied new tools including novel antibodies, biochemical assays, and sequencing approaches to reveal broad expression of the H2B variant H2BE, and defined its role in regulating chromatin structure, neuronal transcription, and mouse behavior. We find that H2BE is enriched at promoters and a single unique amino acid allows it to dramatically enhance chromatin accessibility. Lastly, we show that H2BE is critical for synaptic gene expression and long-term memory. Together, these data reveal a novel mechanism linking histone variants to chromatin regulation, neuronal function, and memory. This work further identifies the first widely expressed H2B variant and uncovers a single histone amino acid with profound effects on genomic structure.

## INTRODUCTION

Histone proteins are the central components of chromatin, affecting DNA compaction and thus, transcription. Two copies of each of the canonical histones—H2A, H2B, H3, and H4—form an octamer around which ∼146 base pairs of DNA are wound^1^. Histones are regulated by post-translational modifications and through exchange with histone variants. Histone variants, which are encoded by separate genes, can substitute for the canonical forms, and are involved in regulation of many cellular processes and gene expression^2^. While canonical histones are transcribed and translated during S phase of the cell cycle and deposited concurrently with DNA replication^3^, histone variants are unique in that they are synthesized throughout the cell cycle and continue to be generated and deposited into chromatin throughout the lifespan of terminally-differentiated cells^4,5^. This feature makes histone variants particularly critical to post-mitotic cells such as neurons^6–15^.

While variants of H2A and H3 are well characterized, there are no known widely expressed H2B variants. One of the few previously discovered H2B variants, H2B.8, is unique to plants and causes chromatin condensation^16^, indicating H2B variants have the potential to affect chromatin structure. A recent study identified seven mammalian H2B variants^17^. Of those, only one known variant, H2BE, is expressed outside of germline tissues. H2BE was discovered in the mouse main olfactory epithelium, where it affects olfactory neuron function^18^. This study concluded that H2BE was only expressed in olfactory epithelial neurons based on transcript expression data. However, due to a lack of a specific antibody, H2BE protein expression could not be analyzed and limited sequencing approaches available at the time precluded analysis of the mechanisms through which H2BE regulates chromatin. While emerging evidence demonstrates the importance of histone variants in the brain and in neurological disorders^6–14^, to the best of our knowledge no subsequent studies have been published on H2BE in the intervening decade. Thus, whether H2BE is expressed outside of the olfactory system and how it affects chromatin and cellular functions beyond olfaction remained a mystery.

Here, we sought to define the function of H2BE in regulating chromatin and examine whether it functions beyond the olfactory epithelium. We generated a highly specific antibody for H2BE, allowing for comprehensive analysis of H2BE. In addition, we combine biochemistry, mouse models, and animal behavior with multiple next-generation sequencing approaches to characterize genomic localization of H2BE as well as its effects on chromatin structure, transcription, and behavior. We show that H2BE is widely expressed throughout the brain. In neurons, H2BE is expressed at gene promoters, conferring an open chromatin conformation. Further, we localize the major effects of H2BE on chromatin to a single amino acid. Loss of H2BE results in widespread chromatin compaction and transcriptional disruption leading to synaptic dysfunction and impaired long-term memory. In summary, this work reveals a novel mechanism by which a histone variant contributes to cognition, and links single amino acid changes in histones to regulation of chromatin structure and transcription.

## RESULTS

### H2BE is enriched at promoters in cortical neurons

The histone variant H2BE differs from its canonical counterpart, H2B, by only five amino acids distributed throughout the length of the protein, with one in each tail and three within the globular domain (Fig 1A). To overcome the lack of tools to specifically detect endogenous H2BE, we developed a highly specific antibody against the first ten amino acids at the N-terminal of H2BE with no cross-reactivity to H2B N-terminal peptides (Fig S1A). To further confirm antibody specificity, we obtained the previously reported H2BE knockout (KO) mouse^18^ and confirmed loss of H2BE transcript in KO mice using RNA-sequencing (Fig S1B-C). As previously reported^18^, KOs are viable and generate expected Mendelian ratios (Fig S1D).

**Figure 1.**
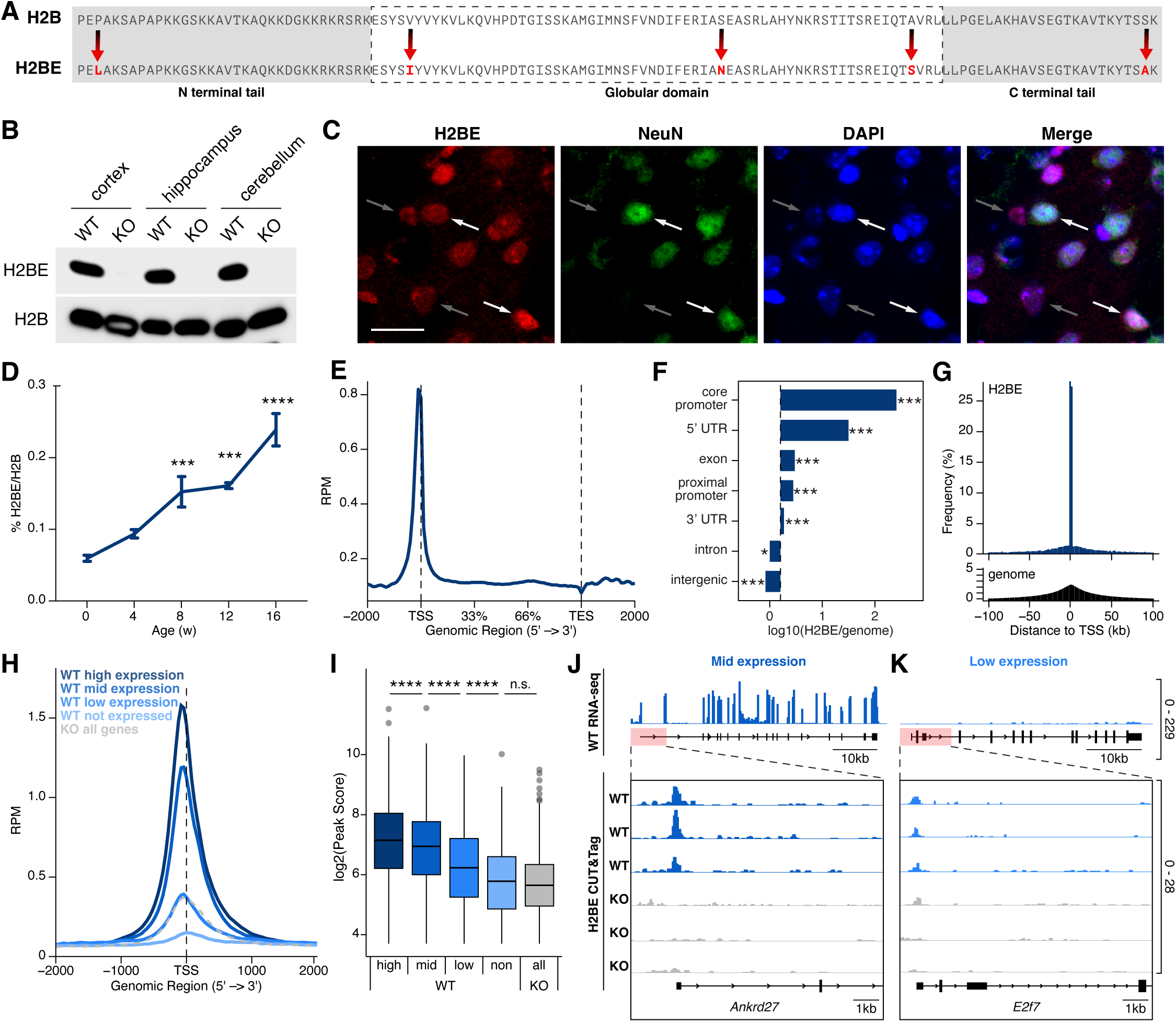
H2BE is enriched at promoters in neurons. (A) H2B and H2BE amino acid sequence. Unique amino acids are highlighted in red. (B) Immunoblot for H2BE and H2B in brain tissue from 3–5-month-old WT and KO mice (ctx = cortex; hpc = hippocampus; crb = cerebellum). H2B serves as loading control. (C) Immunofluorescent images of cortical tissue from 3-month-old mice stained with H2BE, neuronal marker NeuN, and nuclear marker DAPI. Scale bar = 50µm. (D) LC/MS-MS quantification of H2BE in cortical tissue across age (n = 6-10 biological replicates per age group [3-5 male and female combined per age group]; data represents mean +/- SEM; ordinary two-way ANOVA with Dunnett’s multiple comparisons test). No difference was found between male and female mice. (E) Metaplot of H2BE CUT&Tag profiling in WT primary cortical neurons. Plot shows read counts per million mapped reads (RPM) between the transcription start site (TSS) and transcription end site (TES) +/- 2kb (n = 3 biological replicates per genotype). (F) Genomic distribution of H2BE enrichment sites relative to the mouse genome (Chi-square test). (G) Distribution of H2BE enrichment sites (top) and the entire mouse genome binned by 500bp (bottom) relative to the nearest TSS. (H) Metaplot comparison of H2BE CUT&Tag at genes according to expression level. “Not expressed” was defined as genes with mean normalized read counts <3 by RNA-sequencing (n = 8,150 genes). Remaining genes were binned into three equally sized groups by mean normalized read counts (n = 6,028 genes per group). KO shows average across all genes regardless of expression. Plot shows read counts per million mapped reads (RPM) around the TSS +/- 2kb. (n = 3 biological replicates per genotype). (I) H2BE CUT&Tag peak scores assigned by MACS3 by gene expression level as defined above (one-way ANOVA with post-hoc pairwise t-tests with Bonferroni correction). (j-k) RNA-sequencing (top) and CUT&Tag (bottom) gene tracks for representative mid-expressed (*Ankrd27*) and low-expressed (*E2f7*) genes. Statistical details in Table S1. *p<.05,**p<.01,***p<.001,****p<.0001, n.s. = not significant.

We extracted histones from multiple adult mouse tissues and detected widespread expression of H2BE, including in the cortex, hippocampus, and cerebellum of WT mice (Fig S1E, Fig 1B). We found high levels of H2BE protein expression in the main olfactory epithelium, consistent with previous findings^18^. We also measured H2BE in protein lysates from WT cortical tissue and primary cultured cortical neurons with no detectable signal in KOs (Fig S1F-G). We further confirmed specificity using immunohistochemistry of WT and KO brain sections of adult mice (Fig S1H) and found that 64% of H2BE-positive cells co-localize with the neuronal marker NeuN (Fig 1C; Fig S1I), confirming the expression of H2BE in both neurons and nonneuronal cells *in vivo* . These data provide the first characterization of H2BE protein expression, and importantly, demonstrate that it is expressed outside the olfactory system. Further, while H2A and H3 variants are now well characterized in multiple cell types, this analysis of H2BE expression provides the first evidence of a mammalian H2B variant that is widely expressed in multiple tissues.

We next assessed the abundance of H2BE relative to the total pool of H2B in mouse cortex. H2BE was nearly absent in the cortex at birth, while its abundance increased over time and reached 0.24% of all H2B species by adulthood (Fig 1D; Fig S1J). This low percentage is notable given that other histones such as H3.3 and H2A.Z accumulate in neurons and become the dominant species^7,10^. However, histone variants can also have specific genomic enrichment patterns^7,10,19–23^ which may allow H2BE to exert significant effects on chromatin without dominating H2B content. To test this, we performed CUT&Tag with sequencing^24^ on primary cortical neurons from WT and KO E16.5 embryos following 12 days in culture. We confirmed enrichment of H2BE signal over background signal in KO cells (Fig S1K-L) and a high concordance of peaks across biological replicates (Fig S1M). We found that H2BE expression is highly enriched in core promoter regions throughout the genome (Fig 1E-F; Fig S1N) with most identified peaks near a transcription start site (TSS) (Fig 1G) of genes related to important neuronal functions, including dendrite morphogenesis, synaptic vesicle endocytosis, and others (Fig S1O). For further rigor, we validated our CUT&Tag results using H2BE chromatin immunoprecipitation with sequencing (ChIP-seq). Using this orthogonal method of genomic mapping, we found similar enrichment patterns of H2BE (Fig S1P-Q), ensuring that H2BE enrichment around the TSS was not a result of off-target transposase activity at the most open regions of chromatin. For subsequent analysis of H2BE localization, we used CUT&Tag data due to the higher enrichment we observed and lower read depth required to achieve high-confidence peak calls. Further, H2BE enrichment is positively correlated with gene expression in WT neurons (Fig 1H-K), suggesting a relationship between H2BE enrichment and gene expression. Together, these data indicate that H2BE is highly enriched at core promoters, a region critical for the control of transcription.

### H2BE promotes open chromatin via a single amino acid

H2BE has 3 unique amino acids that lie within the globular domain. Of these, N75 and S97 are embedded within the nucleosome core, while I39 lies within the histone-DNA interface^1^ (Fig 1A). Due to the location of 2 amino acids within the nucleosome core, we tested whether H2BE affects nucleosome stability. We generated nucleosomes containing either two copies of canonical H2B or variant H2BE with canonical H2A, H3, and H4. Importantly, there were no changes in thermal stability between H2B- and H2BE-containing nucleosomes. Both species yielded two melting peaks at ∼72°C (Tm1) and ∼85°C (Tm2) corresponding to H2A/H2B dimer and H3/H4 tetramer dissociation, respectively (Fig 2A). This indicates that H2BE-specific amino acids do not affect nucleosome stability.

**Figure 2.**
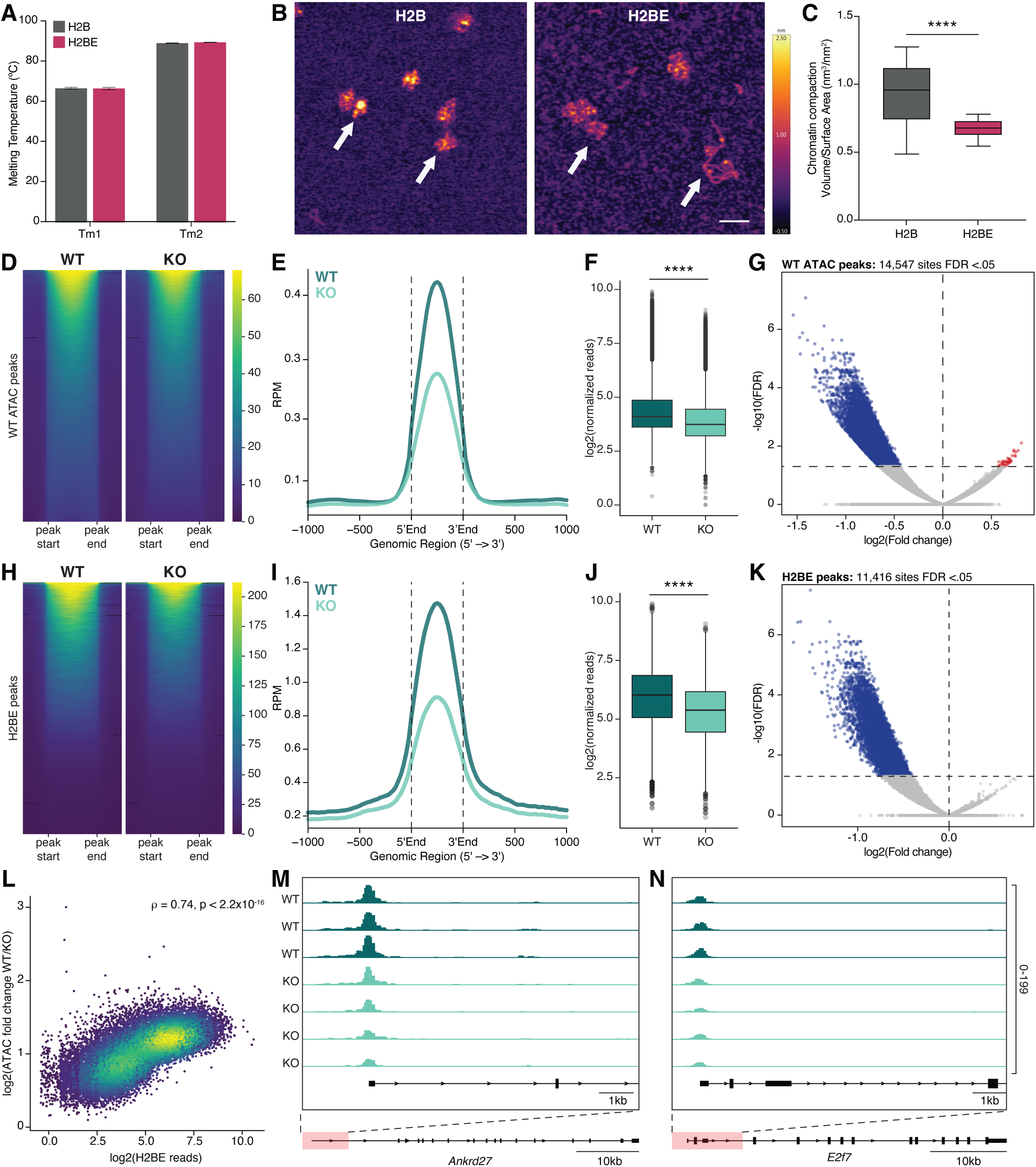
H2BE promotes open chromatin. (A) Thermal stability assay of H2B- and H2BE-containing nucleosomes yields two melting peaks (Tm) corresponding to dimer and tetramer dissociation, respectively (n = 3 replicates per group; unpaired Welch’s t-tests with Holm-Šídák correction). (B) Representative micrographs of H2B- or H2BE-containing nucleosome arrays. Scale bar = 100µm. (C) Volume/surface area quantification of compaction from atomic force microscopy of H2B or H2BE nucleosome arrays (n = 60 H2B arrays, 40 H2BE arrays; Welch’s t-test). Surface area = Total # of pixels occupied by array. Volume = Total pixel intensity of array. (D) Heat map and (E) metaplot comparison of ATAC-seq average signal from WT and KO primary cortical neurons at all WT peaks. Plot shows read counts per million mapped reads (RPM) at all measured peaks +/- 1kb (n = 3 WT biological replicates, 4 KO biological replicates). (F) Normalized ATAC-seq read counts at all ATAC peak sites detected in WT neurons (unpaired t-test). (G) Volcano plot showing differential chromatin accessibility between WT and KO at WT ATAC peaks. Blue = peaks that decrease in KO; red = peaks that increase in KO; gray = peaks below significance cut-off of FDR<.05. (H) Heat map and (I) metaplot comparison of ATAC-seq average signal from WT and KO primary cortical neurons at all H2BE binding sites identified by CUT&Tag. Plot shows read counts per million mapped reads (RPM) at all measured peaks +/- 1kb (n = 3 WT biological replicates, 4 KO biological replicates). (J) Normalized ATAC-seq read counts at H2BE binding sites (unpaired t-test) (K) Volcano plot showing differential chromatin accessibility at H2BE binding sites. Blue = peaks that decrease in KO; red = peaks that increase in KO; gray = peaks below significance cut-off of FDR<.05. (L) Correlation between H2BE reads in WT neurons and changes in ATAC-seq signal between WT and KO (Spearman correlation; ρ = 0.74; p < 2.2×10^-16^). (M-N) ATAC-seq gene tracks for representative mid-expressed (*Ankrd27*) and low-expressed (*E2f7*) genes. ****p<.0001.

We next tested the ability of H2BE to compact DNA. Nucleosome arrays were generated by combining H2B- or H2BE-containing octamers with 12×601 DNA and compaction was measured using atomic force microscopy^25^. We found that canonical H2B-containing arrays occupy a smaller surface area compared to H2BE-containing arrays (Fig 2B-C), without a change in volume of the nucleosomes (Fig S2A), demonstrating that H2BE confers a more open chromatin conformation relative to canonical H2B. To confirm these findings, we used Mg^2+^ precipitation assays^26^ and found that H2BE arrays display a lower propensity to oligomerize in the presence of increasing concentrations of Mg^2+^ (Fig S2B). Together, these assays revealed that H2BE nucleosome arrays are more open than H2B arrays without affecting nucleosome stability. Further, these data indicate that this effect is an innate property of H2BE and intrinsic to the amino acid sequence, as the only components present in these *in vitro* assays include unmodified histones and DNA without other protein complexes or histone modifications.

These biochemical assays were performed using synthetic nucleosomes containing two copies of either H2B or H2BE. However, in a cellular context, it is possible that H2BE coexists with H2B in a nucleosome. We therefore assessed the effect of H2BE on chromatin accessibility in a physiological context using WT and KO primary cortical neurons paired with the assay for transposase-accessible chromatin with sequencing (ATAC-seq)^27^. We first examined the relationship between WT chromatin accessibility and H2BE enrichment and found a strong positive correlation throughout the genome (Fig S2C). We next performed differential analysis of chromatin accessibility in KO at accessible regions including transcription start sites (TSSs) in an unbiased fashion regardless of whether a peak was detected in WT neurons. We found a decrease in accessibility in KO neurons at TSSs (Fig S2D-F) with 99.98% of differential sites (8,559) downregulated, while only 0.02% (2 sites) were upregulated. We observed a similar decrease at enhancers (Fig S2G-I), with no sites showing significantly increased accessibility. We also examined all peaks detected in WT neurons regardless of genomic region, and again found that loss of H2BE results in markedly decreased chromatin accessibility (Fig 2D-G; Fig S2J-K).

To determine the relationship between H2BE enrichment and chromatin accessibility, we measured ATAC-seq signal at H2BE peaks identified by CUT&Tag. Of all H2BE peaks, 53.2% had decreased accessibility in KO, and zero had increased accessibility (Fig 2H-K). Further, we found that sites with greater enrichment of H2BE in WT neurons showed greater decreases in accessibility, suggesting a direct correlation between H2BE occupancy and chromatin compaction upon H2BE loss (Fig 2L-N). H2BE-bound transcription start sites showed the greatest decreased accessibility, while more modest changes occurred in gene body and intergenic H2BE peaks (Fig S2L-M). To better understand the precise location of H2BE in relation to the TSS, we examined ATAC-seq and H2BE CUT&Tag peaks within TSS regions. We measured the distance of peak summits to the TSS and found that H2BE peaks are further upstream from the TSS compared to ATAC-seq peaks, suggesting enrichment at the -1 nucleosome (Fig S2N). These data demonstrate a direct role of H2BE in modulating chromatin accessibility in neurons.

To identify the site driving the effects of H2BE on chromatin accessibility, we focused on I39 based on its location at the nucleosome-DNA interface (Fig 3A) and its conservation in human H2BE. We generated mutant nucleosomes in which the H2BE-specific I39 is mutated to valine (H2BE-I39V) found on canonical H2B, and the converse mutation for canonical H2B (H2B-V39I) (Fig 3B). We detected no differences in thermal stability of the mutant nucleosomes (Fig 3C). We then constructed chromatin arrays from these nucleosomes (Fig S3A-B) and measured compaction with a Mg^2+^ precipitation assay. We found that arrays containing H2BE-I39V are less soluble than WT H2BE arrays in the presence of Mg^2+^ (Fig 3D). Further, the V39I mutation on canonical H2B was sufficient to increase solubility (Fig 3E). Together, these data demonstrate that I39 on H2BE is both necessary and sufficient for H2BE to open chromatin. Finally, we generated viral vectors containing WT H2BE or mutant H2BE-I39V to re-express H2BE in KO neurons (Fig S3C-D). Using ATAC-seq we again found that WT neurons have more open chromatin than KO neurons and that re-expressing WT H2BE was sufficient to open chromatin around the TSS. We confirmed that these results were not due to viral infection or overexpression of a histone using a virus expressing GFP or canonical H2B, respectively. Further, neurons expressing I39V mutants were indistinguishable from KO neurons, confirming the necessity of this amino acid to open chromatin in a cellular context (Fig 3F-G). To determine if mutating any of the 5 H2BE-specific amino acids affects H2BE function, we performed ATAC-seq on KO neurons expressing mutant H2BE in which L3 on H2BE was mutated to proline (L3P) as found in canonical H2B (Fig S3D-E). Because this amino acid is found in the N-terminal tail of H2BE, we predicted it would have no role in chromatin compaction. Notably, H2BE-L3P had the same effect on chromatin compaction as WT H2BE (Fig S3F). Taken together, these data illustrate that I39 is necessary for the effect of H2BE on chromatin compaction.

**Figure 3.**
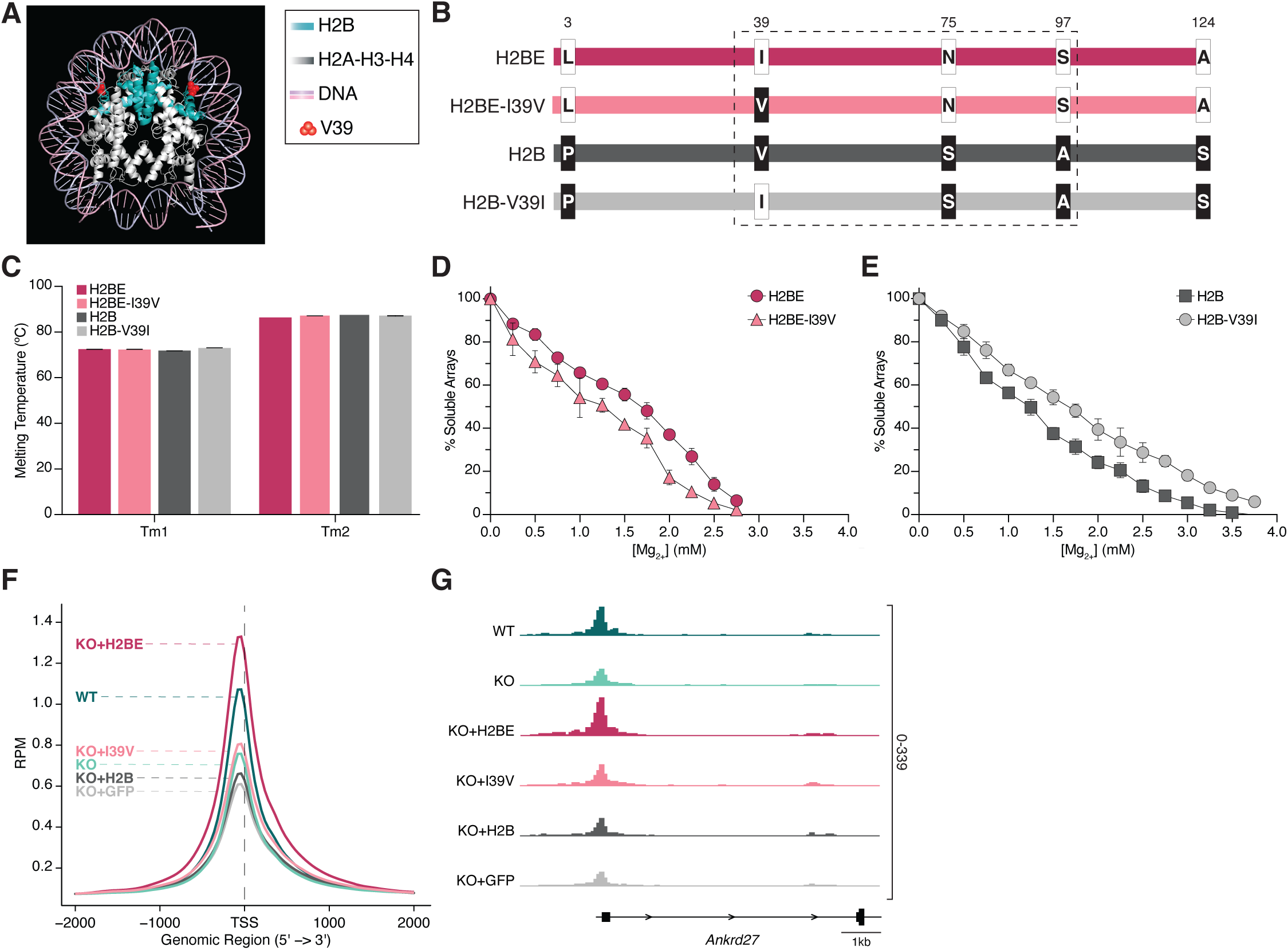
Single amino acid at the histone-DNA interface drives H2BE effect on chromatin. (A) Nucleosome surrounded by DNA shows that the H2BE-specific amino acid at site 39 lies at the histone-DNA binding interface. Cyan=H2B; red=V39; white=H2A/H3/H4. (B) Schematic of canonical H2B, WT H2BE, and mutant H2B/H2BE sequences. (C) Thermal stability assay of H2BE and H2BE-I39V nucleosomes (n = 3 biological replicates per group; 2-way ANOVA). (D) Mg^2+^ precipitation assay of chromatin arrays containing WT H2BE or H2BE-I39V (n=4 per group) (E) Mg^2+^ precipitation assay of chromatin arrays containing canonical H2B or H2B-V39I (n=8 per group) (F) ATAC-seq average signal at all genes (n = 3-8 biological replicates per condition). Plot shows read counts per million mapped reads (RPM) around the transcription start site (TSS) +/- 2kb. (G) ATAC-seq gene tracks for *Ankrd27*. Statistical details in Table S1.

### H2BE promotes synaptic gene expression

Given the profound changes in chromatin accessibility in KO neurons, we next performed RNA-sequencing on WT and KO primary cortical neurons to determine how loss of H2BE affects gene expression. We saw broad changes in gene expression in KO neurons, with 1,790 downregulated and 1,764 upregulated genes (Fig 4A). We note that *H2bc21* is called as upregulated in KO in our differential expression analysis because of reads mapping to the 3’UTR located downstream of the H2BE coding sequence (Fig S1B-C) which is replaced by mCherry^18^, allowing for continued expression of the UTR. Downregulated genes are enriched for functions related to neuronal morphology and axon growth while upregulated genes are enriched for cellular metabolism and mitochondrial function (Fig 4B). Further analysis with SynGO^28^ confirmed disruption of critical synaptic genes related to regulation of synaptic transmission and signaling (Fig S4A).

**Figure 4.**
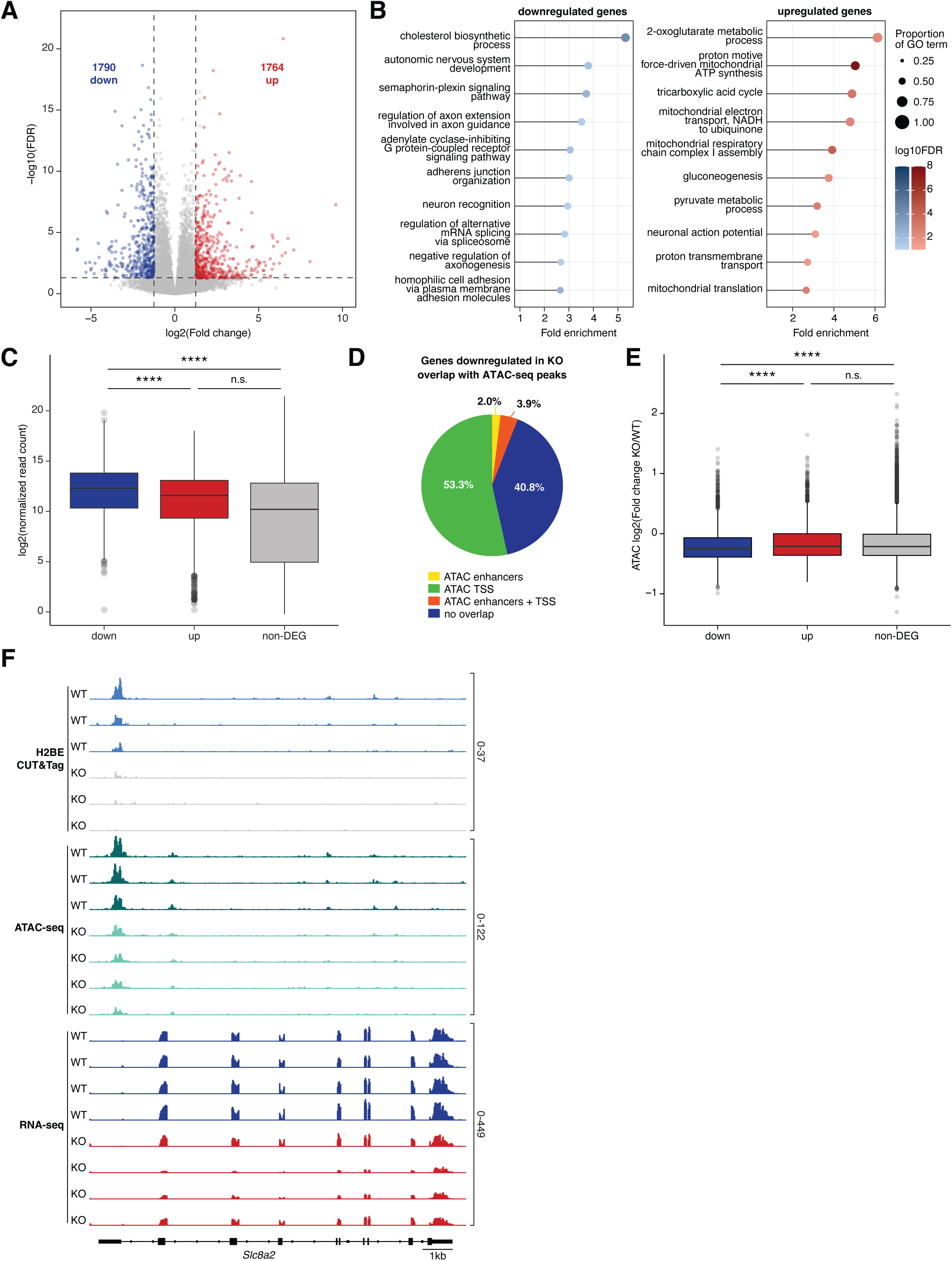
H2BE promotes synaptic gene expression. (A) Volcano plot showing differentially expressed genes (FDR < .05, absolute fold change > 1.25) between WT and KO primary cortical neurons (n = 4 biological replicates per genotype [2 male, 2 female]). Blue = downregulated; red = upregulated. (B) Gene ontology enrichment analysis of down-(left) and upregulated (right) genes. (C) Normalized RNA-seq read counts in WT neurons for genes downregulated, upregulated or unchanged in KO (one-way ANOVA with post-hoc pairwise t-tests with Bonferroni correction). (D) Pie chart showing which downregulated genes have a decreased ATAC-seq peak in KO at a proximal enhancer and/or promoter for that gene in WT. (E) ATAC-seq fold change (KO/WT) for downregulated, upregulated, and unchanged genes in KO (one-way ANOVA with post-hoc pairwise t-tests with Bonferroni correction). (F) CUT&Tag, ATAC-seq, and RNA-seq genome browser tracks at the *Slc8a2* locus in WT and KO cortical neurons. See Fig S4 for additional example tracks. Statistical details in Table S1. RNA-seq gene lists and full GO in Table S2. ****p<.0001, n.s. = not significant; DEG = differentially expressed genes.

We found that genes downregulated in KO had higher expression in WT cells compared to both upregulated and non-differentially expressed genes (Fig 4C), suggesting that highly expressed genes are more sensitive to H2BE loss. We used ATAC-seq data to determine the relationship between changes in accessibility and gene expression and found that ∼60% of downregulated genes show decreased accessibility at the TSS and/or proximal enhancers (Fig 4D). Further, downregulated genes have the greatest decrease in accessibility (Fig 4E). Taken together, these data suggest that genes downregulated in KO are the most highly expressed in WT and show the greatest changes in accessibility upon H2BE loss. Downregulated genes without differential ATAC peaks and upregulated genes may be due to secondary or compensatory mechanisms or other yet unknown functions of H2BE. Finally, overlap of CUT&Tag, ATAC, and RNA sequencing datasets identified genes with H2BE enrichment in WT neurons, changes in chromatin compaction, and decreased transcription in KO, including *Slc8a2 and Tuba1a* (Fig 4F; Fig S4B-E).

### H2BE affects neuronal gene expression in the brain

We next sought to understand the role of H2BE on gene expression the brain. Given that H2BE is expressed at negligible levels early in life, we predicted H2BE loss would not have substantial effects on cell identity but may affect gene expression within cell types due the cumulative effects of H2BE loss throughout adulthood. We performed single-nucleus Drop-seq^29^ on cortical tissue from adult male WT and KO mice. We identified 20 unique cell clusters comprising all expected cell types without notable changes in cluster identity across genotypes (Fig 5A-B, Fig S5A). Cell identity was determined using well-established marker genes for major cortical cell types (Fig S5B)^29–33^. Specifically, we identified 10 excitatory neuronal clusters (*Slc17a7*^+^), 5 inhibitory neuronal clusters (*Gad2*^+^), oligodendrocytes (*Mog*^+^, *Enpp6*^+^, *Opalin*^+^), microglia (*Ctss*^+^) and endothelial cells (*Flt1*^+^). We then used known cortical layer-specific markers to reveal excitatory neuronal subtypes (Layer 2/3: *Ndst4*^+^, *Enpp2*^+^; Layer 4: *Rorb*^+^; Layer 5: *Tox*^+^; Layer 6: *Zfpm2*^+^, *Hs3st4*^+^; Layer 6b: *Ctgf*^+^). We identified differentially expressed genes in almost all neuronal clusters (Fig 5C; Fig S5C), and in non-neuronal cells, including astrocytes, oligodendrocytes, and microglia (Fig S5C).

**Figure 5.**
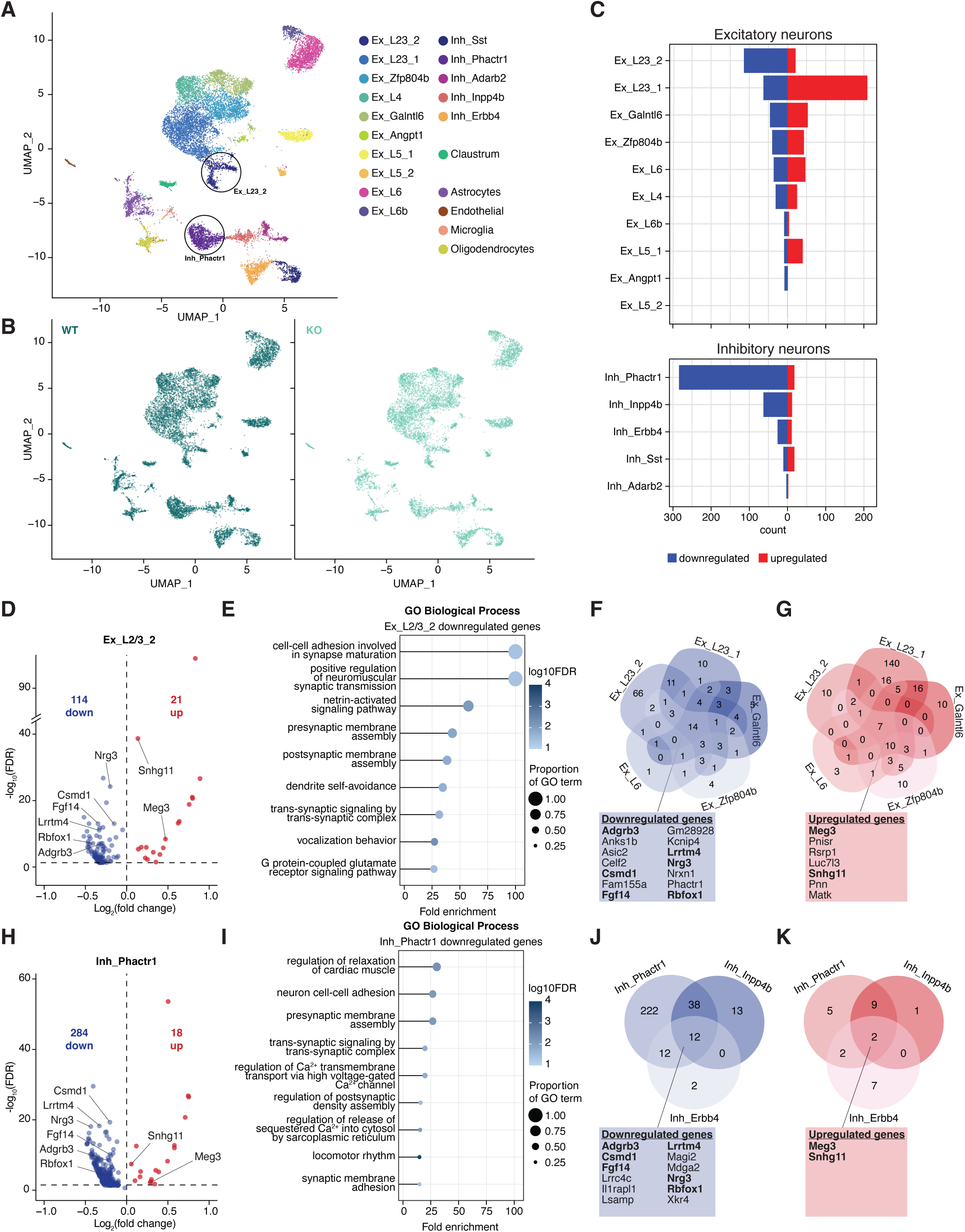
H2BE is required for appropriate synaptic function in the brain. (A) UMAP of single-nucleus transcriptomic profiles from adult male (2-4 months) mouse cortices (n = 3 biological replicates per genotype). Colors represent cluster identity. (B) UMAP clustering with dots representing each nucleus colored by genotype. Relative distribution of WT and KO cells found by cluster shown in Figure S5. (C) Number of differentially expressed genes within each cluster for all excitatory and inhibitory neuronal clusters. (D) Volcano plot showing differential expression from WT and KO in a representative cluster of layer 2/3 cortical excitatory neurons (Ex_L2/3_2). (E) Gene ontology enrichment analysis of downregulated genes in cluster Ex_L2/3_2. Upregulated shown in Figure S5. (F-G) Overlap of differentially expressed down-(F) and upregulated (G) genes from the 5 excitatory neuron clusters with the greatest number of differentially expressed genes. (H) Volcano plot showing differential expression from WT and KO cortical inhibitory neurons in a representative cluster, Inh_Phactr1. (I) Gene ontology enrichment analysis of downregulated genes in cluster Inh_Phactr1. (J-K) Overlap of differentially expressed down-(J) and upregulated (K) genes from the 3 inhibitory neuron clusters with the greatest number of differently expressed genes. Ex = excitatory neurons; Inh = inhibitory neurons; blue = downregulated genes; red = upregulated genes.

Of all excitatory neurons, the two clusters from cortical layer 2/3 had the most differential gene expression. Looking at the excitatory cluster with the greatest number of downregulated genes from layer 2/3 pyramidal neurons (Ex_L2/3_2), we observed more downregulated genes (114) than upregulated (21), a common feature of most cell types (Fig 5D-E; Fig S5D). These downregulated genes are involved in synaptic functions such as cell-cell adhesion (*Nlgn1, Nrxn1* ), pre- and postsynaptic membrane assembly (*Il1rapl1, Magi2* ), and trans-synaptic signaling (*Grm7, Tenm2*) (Fig 5E). Overlapping the down- and upregulated genes from the top 5 excitatory clusters identified multiple commonly disrupted genes (Fig 5F-G). The top inhibitory cluster also showed mainly downregulated genes with related functions to those from layer 2/3 pyramidal neurons (Ex_L2/3_2) (Fig 5H-I; Fig S5E). We also identified genes disrupted across multiple inhibitory clusters and genes common between all top excitatory and inhibitory clusters (Fig 5J-K). In summary, H2BE is essential for the expression of synaptic genes across multiple cell types in the adult mouse brain.

### H2BE promotes synaptic strength and long-term memory

These intersecting changes in synaptic gene expression raise the possibility that knocking out H2BE will lead to weakened synapses. To assess synaptic function, we performed field recordings of pharmacologically isolated excitatory postsynaptic potentials (fEPSP) from the stratum radiatum of the CA1 region in acute hippocampal slices. We electrically stimulated Schaffer collateral fibers with increasing input voltage, observing increased slope of the resulting fEPSP. However, the response is significantly lower in KO, suggesting that synapses are weaker in KO in response to the same intensity stimulus (Fig 6A). To rule out that differences in synaptic strength resulted from altered recruitment of Schaffer collaterals, we measured fiber volley amplitude and saw no difference between WT and KO (Fig 6B-C). These data suggest that excitatory glutamatergic synaptic transmission is weakened in KO, in line with our finding that H2BE promotes the expression of synaptic genes.

**Figure 6.**
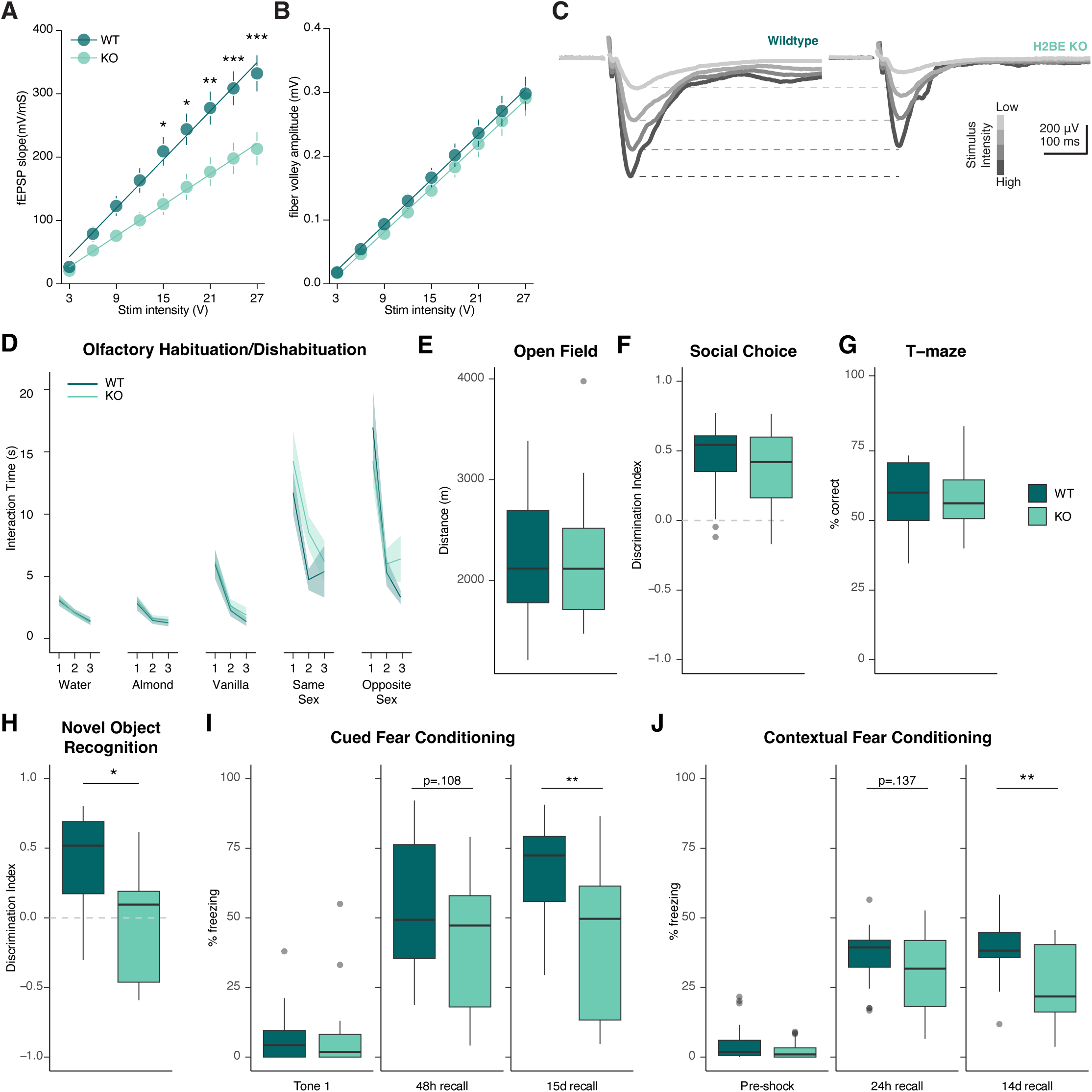
H2BE promotes synaptic strength and long-term memory. (A) Input-output curves from field extracellular postsynaptic potentials and (B) afferent fiber volleys following stimulation of hippocampal Schaffer collaterals. The line represents the inferred responses through linear regression (WT r^2^=0.6748, KO r^2^=0.5006), while each dot corresponds to the averaged slope from each experiment based on stimulus intensity. (n = 11/13 from 5/4 animals [2-3 males and females per genotype]; two-way ANOVA [stimulus intensity x genotype] with Šídák’s multiple comparisons test). (C) Representative traces from WT and KO. (D) Interaction time with a scented cotton swab during presentations of water, two non-social odors (almond, vanilla), and social odors (same sex, opposite sex) for three consecutive trials per scent. Line and shaded area represent mean +/- SEM (n = 20 WT mice [8 male, 12 female], 24 KO mice [10 male, 14 female]; three-way ANOVA [genotype x sex x trial number] for each odorant. No differences were detected between genotypes. Significant differences were detected between sex (graphed separately in Figure S6A). (E) Distance traveled in an open field arena during a 5-minute trial (n = 19 WT mice [9 male, 10 female], 19 KO mice [11 male, 8 female]; two-way ANOVA [genotype x sex]). Sexes graphed separately in Figure S6B. (F) Discrimination index for interaction time between a mouse and a rock during a 3-chamber sociability SEM (n = 20 WT mice [8 male, 12 female], 24 KO mice [10 male, 14 female]; two-way ANOVA [genotype x sex]). DI = ([mouse – rock] / [mouse + rock]). No effects of sex were observed. (G) Percent correct trials of spontaneous alternation in a T-maze trial (n = 19 WT mice [9 male, 10 female], 19 KO mice [11 male, 8 female]; two-way ANOVA [genotype x sex]). No effects of sex were observed. (H) Discrimination index during a novel object recognition task (n = 19 WT mice [9 male, 10 female], 19 KO mice [11 male, 8 female]; two-way ANOVA [genotype x sex]; *p<.05)]). DI = ([novel – familiar] / [novel + familiar]). No effects of sex were observed. (I-J) Percent freezing during cued (I) and contextual (J) fear conditioning (n = 20 WT mice [8 male, 12 female], 24 KO mice [10 male, 14 female]; two-way ANOVA [genotype x sex] for each phase of testing). Statistical details in Table S1. *p<.05,**p<.01,***p<.001.

Alterations in synaptic function can cause deficits in cognition. Therefore, we performed several behavioral assays to analyze the role of H2BE in cognition and learning. First, we assessed olfactory function due to the high level of H2BE expression in mouse olfactory tissues^18^. Previous work found that H2BE-KO caused a deficit in olfactory discrimination in a learning paradigm. Based on findings that H2BE is expressed throughout the brain (Fig 1B, Fig S1E-G), it is unclear if these deficits are due to olfactory dysfunction or indicative of broader learning and memory deficits. We therefore first tested olfactory habituation and dishabituation^34^ through repeated exposure to the same odorant followed by exposure to novel odorants. We found that KO mice have intact habituation and dishabituation to a variety of non-social and social olfactory cues (Fig 6D). There was a significant effect of sex on time spent interacting with the olfactory cues, with females showing greater interaction time with odor cues but no effect of genotype (Fig S6A). Notably, these findings fit with published data demonstrating that KO mice have intact odor evoked electrical responses in the olfactory epithelium^18^ but raise the possibility that deficits may be due to other cognitive disruptions. In an open field arena, WT and KO mice traveled the same distance when allowed to explore freely, with male mice traveling a shorter overall distance compared to female mice (Fig 6E; Fig S6B). There was no effect of H2BE-KO on anxiety-like behavior as assessed by time spent in the center zone of an open field arena (Fig S6C). These behavioral tests indicate KO mice have intact olfaction and no detectable mobility deficits or changes in anxiety and exploratory behaviors.

Due to the well-documented role of synaptic genes in social behaviors^35–37^, we tested the sociability of WT and KO mice. Mice were placed in a three-chamber apparatus containing a rock or mouse in each of the outer chambers and a neutral center chamber. Mice were allowed to explore freely, and we measured the amount of time spent interacting with the mouse and rock. A discrimination index was calculated for each mouse by finding the difference between interaction time with the mouse and rock as a proportion of total time spent interacting with both. Both WT and KO mice show a preference for the mouse (as measured by a positive discrimination index) (Fig 6F). However, subtle changes were detected in average interaction duration with the mouse (Fig S6D-F), suggesting largely intact sociability with minor changes in social behavior.

We next tested KO and WT littermates in a T-maze to test working memory. WT and KO mice exhibited similar rates of 3-arm alternations with no effect of sex (Fig 6G), suggesting that exploratory behavior and working memory remain intact in KO mice. Finally, we tested long-term memory, selecting tests that rely on proper synapse regulation^38–40^. In a novel object recognition test, mice are exposed to two identical objects in an arena and allowed to explore freely. On the following day, one of the objects is replaced with a novel object. We observed that mice lacking H2BE were not able to distinguish between a novel object and a familiar object presented 24 hours earlier with no effect of sex (Fig 6H), suggesting that long-term memory is disrupted in KO mice. Importantly, there were no differences between groups in total time spent with both objects during learning or testing (Fig S6G-H), indicating that deficits were not due to differences in object exploration.

We also assessed mice in two complementary tests of long-term memory—cued and contextual fear conditioning with two foot-shocks paired with a tone. In contextual conditioning trials, mice are returned to the chamber or “context” while for cued conditioning trials, mice are placed in an altered context and re-exposed to the tone cue alone (no foot-shock). In both assays, freezing is measured as a fear response. There was no difference in freezing behavior prior to the initial tone-shock pair (“Pre-shock”) or in response to the first tone presentation (“Tone 1”), illustrating that there was nothing inherently fearful about the context or tone cue alone. No effect of sex was observed in any fear conditioning assays. However, KO mice exhibit less freezing behavior 14 (or 15) days post-training compared to their control littermates (Fig 6I-J). Interestingly, in both contextual and cued conditioning, knockout mice showed a trend toward decreased freezing at 24 hour (contextual) or 48 hour (cued) post-training, however this difference did not reach significance. Together, these results demonstrate a critical role for H2BE in long-term memory, while olfaction, motor behavior, exploration, social behavior, and working memory are largely spared.

In summary, these findings support an overarching model in which H2BE promotes accessibility at critical and highly expressed neuronal genes to support synaptic responses and robust memory. Thus, when H2BE is lost, chromatin adopts a more closed state and synaptic gene expression is reduced, leading to synaptic dysfunction and impaired long-term memory (Fig 7).

**Figure 7.**
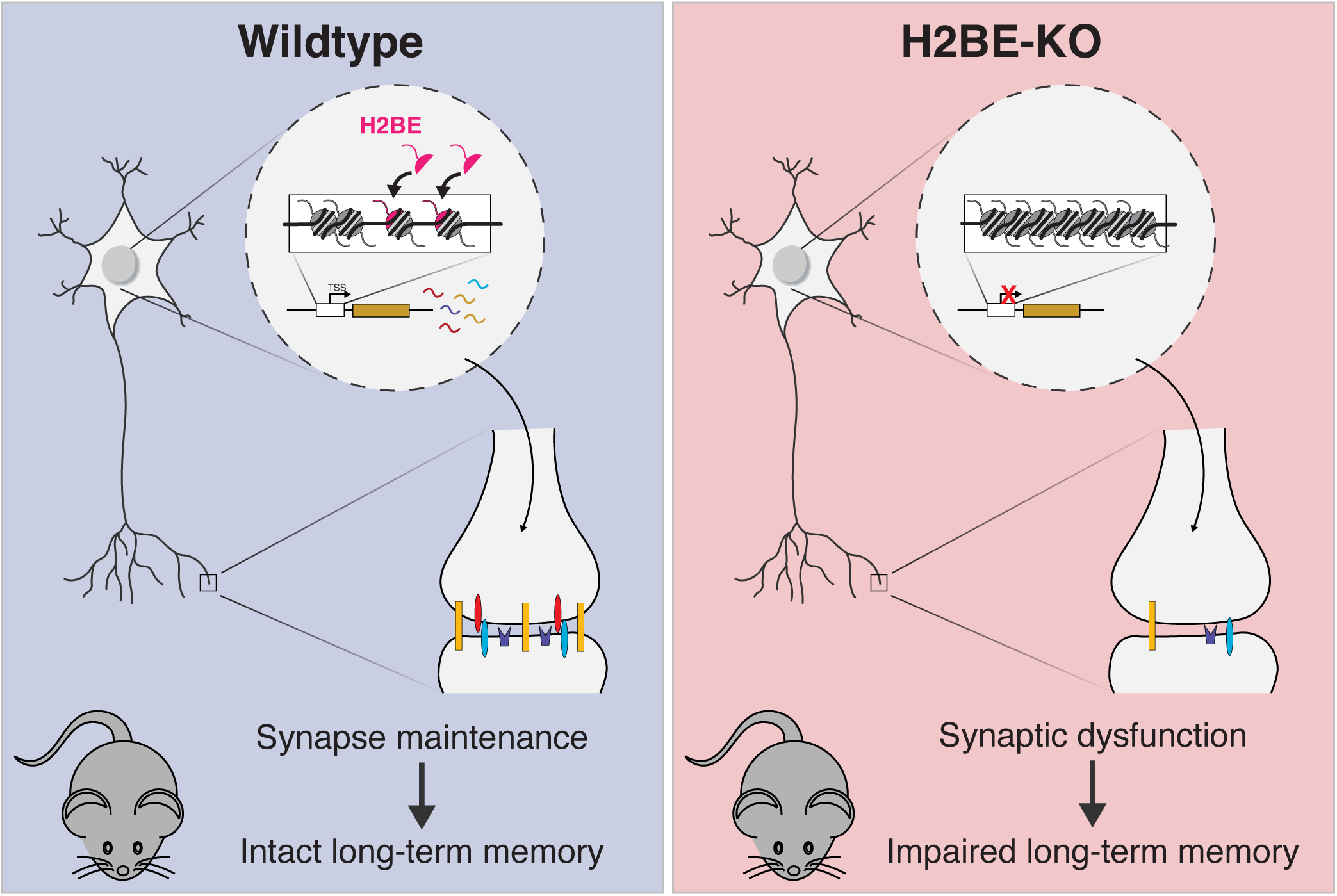
Model of H2BE function. In a WT mouse, H2BE is expressed at promoters throughout the genome, where its expression causes chromatin to open and higher levels of synaptic gene expressions. This supports healthy synapses and proper neuronal function. In contrast, upon loss of H2BE chromatin remains in a closed state, there is loss of synaptic gene expression, synaptic dysfunction, and impaired long-term memory.

## DISCUSSION

Here, we show widespread expression of H2BE in mice, with enrichment at gene promoters in the brain. The expression of H2BE results in greater chromatin accessibility which is reliant on a single amino acid that lies at the interface between the histone octamer and nucleosomal DNA. Further, we show that loss of H2BE results in broad changes in gene expression in both primary neuronal cultures and in neurons derived from the adult brain. Lastly, we show that H2BE is critical for long-term memory. In summary, we provide evidence of a widely expressed H2B variant with an innate ability to control chromatin structure and a novel example of how histone variants affect chromatin structure, transcription, and memory.

Of the five amino acids that differ in H2BE compared to canonical H2B, we found that one is critical for its effects on chromatin accessibility. Our data thus place H2BE as a novel example where a single amino acid on a histone is linked to large-scale changes in chromatin regulation. This has recently been shown in two other variants, H2A.Z and H3.3, where a single amino acid is linked to neurological function and the onset of neurodevelopmental disorders^14,20^. Additionally, emerging research on oncohistones reveals how single amino acid mutations in histone variants cause genome-wide changes underlying various cancers^41–44^. Our work adds to this emerging theme in chromatin biology and provides a novel example of a variant-specific amino acid that mediates robust changes in chromatin structure.

This study also defines H2BE as the first mammalian H2B variant with widespread expression. In line with previous work^18^, we see substantial enrichment of H2BE in the mouse MOE (Fig S1E). However, the development of an H2BE-specific antibody, H2BE nucleosomes and chromatin arrays, along with the expansion of next-generation sequencing technologies has allowed us to fully define H2BE expression and its function in chromatin regulation. Notably, none of our findings disagree with previous work but use newly available tools to provide an in-depth analysis of H2BE expression and function. Santoro et. al. discovered exceptionally high levels of H2BE in olfactory neurons of the mouse main olfactory epithelium^18^. Notably, these neurons are quite distinct from cortical neurons in that they exhibit a remarkably high amount of neuronal turnover compared to the rest of the nervous system. In contrast, we found relatively low levels of H2BE in the cortex with H2BE making up <1% of the total pool of H2B. Direct comparison of MOE, cortex, and other tissues further supports high levels in the MOE compared to other tissues (Fig S1E). We expect that this drastic difference in expression likely explains the different outcomes of H2BE function in these distinct regions. Our biochemical results demonstrate that the ability to open chromatin is an innate feature of H2BE that does not require cellular components and thus is expected to occur at sites of H2BE incorporation regardless of cell type or tissue. Replacing the majority of H2B with H2BE—as likely occurs in inactive neurons in the MOE—could therefore be expected to cause a drastic decompaction of the chromatin fiber resulting in cell death. Interestingly, loss of compaction can cause genomic instability and aberrant transcriptional de-repression, including in the mouse brain^45–47^. In addition, global loss of canonical histones and widespread heterochromatin loss is a hallmark feature of aging and cell loss^45,48–50^. While outside the scope of this study, it is possible that olfactory neurons co-opted such a mechanism to control cell fate in response to low olfactory cues. However, we do not anticipate that a similar mechanism is likely to occur in a healthy cortex where high levels of cell death and neuronal turnover does not occur. Rather, in the cortex, H2BE is restricted to a small percentage of the total pool of H2B and our data suggest neurons use it to open chromatin at precise genomic sites where high gene expression is required.

Interestingly, there is a growing body of clinical data linking histone variants to neurological disorders. H2BE-KO mice have memory disruptions but no overt health issues or disruptions in basic motor and cognitive functions, suggesting that the cognitive deficits that we observed are not due to major health issues. However, it is plausible that H2BE has distinct roles in other cell types and we detected H2BE in non-neuronal cell types in the brain. Further, the role of H2BE in other cell types could emerge in other biological contexts, including disease, aging, or other perturbations. To date H2BE has not been linked to neurological or neurodevelopmental disorders. However, a recent study of human histone genes suggests that H2BE is intolerant to mutations, suggesting that mutations in H2BE may be pathogenic^51^. We anticipate that with increased use of whole exome sequencing in patients with unexplained neurodevelopmental disorders, it is likely that future work will reveal mutations within H2BE, similar to disorders recently linked to other histone variant proteins^6,20^.

In summary, this work reveals how histone variant H2BE contributes to the complex chromatin environment in the brain. This research furthers our understanding of mechanisms by which neurons control transcription and ultimately govern behavior. Further, this work provides insight into novel mechanisms underlying memory and the increasing number of neurodevelopmental disorders linked to disruptions in chromatin regulation.

## LIMITATIONS OF THE STUDY

With the finding that H2BE is expressed throughout the brain and controls chromatin accessibility, several questions remain. The mechanism through which H2BE is enriched at promoters and whether it has additional effects on chromatin remain unknown. Specific chaperone proteins may deposit H2BE as occurs for other variants. Alternatively, greater histone turnover at regions of open chromatin may result in the accumulation of H2BE at those sites which then serves to maintain accessible chromatin states. Further, while we observe large-scale changes in transcription upon H2BE-KO, ∼60% of downregulated genes can be explained by a loss of accessibility. We speculate that other differentially expressed genes may be explained by secondary or compensatory mechanisms, or by functions of H2BE that have not yet been discovered.

The mechanisms governing H2BE expression also remain unclear. Notably, like other histone variants H2BE accumulates with age. This accumulation is presumably a consequence of the extended time in which neurons exist in a postmitotic state when they can no longer generate large quantities of replication-dependent histones but continue to generate variants. Whether an increase in H2BE content in neurons is balanced by other chromatin regulatory mechanisms or is either detrimental or beneficial to cell health in the aging brain remains unclear. In addition, previous work on H2BE in olfactory neurons of the mouse main olfactory epithelium found that H2BE is inversely correlated to neuronal activity. Here we focus on the function of H2BE under basal conditions and proposed a model in which H2BE sustains high expression of critical synaptic genes to maintain an appropriate transcriptional state. However, our model does not define the effect of neuronal activity on expression of H2BE. We expect that pursuing these lines of inquiry would provide critical insights into H2BE function across brain regions, cell types, and lifespan.

## Acknowledgements

We thank Drs. Catherine Dulac and Stephen Santoro for sharing reagents and mouse lines. Mass spectrometry was performed with the CHOP-Penn Proteomics Core. Behavior procedures were performed with assistance from The Neurobehavior Testing Core at UPenn/ITMAT and IDDRC at CHOP/Penn U54 HD086984. We acknowledge the use of the MSK Molecular Cytology Core Facility, supported by the NCI Cancer Center Support Grant (CCSG, P30 CA08748), Cycle for Survival, and the Marie-Josée and Henry R. Kravis Center for Molecular Oncology. E.H. is supported by NIH grant F31MH126576. S.L, K.P., and E.K., were supported by NIH grants 1DP2MH129985 and R00MH111836 and by the Klingenstein-Simons Fellowship from the Esther A. & Joseph Klingenstein Fund and the Simons Foundation, the Alfred P. Sloan Foundation Research Fellowship (FG-2020-13529), the Brain and Behavior Research Foundation NARSAD Young Investigator Award, and pilot funding from the Epigenetics Institute at the University of Pennsylvania. N.A.P. is supported by NIH grant F99CA264420.

## Author Contributions

E.R.F. designed, performed and analyzed the results for most experiments, and wrote the manuscript. S.L. generated key reagents and provided input and expertise. N.A.P., T.B., and Q.G. performed and analyzed biochemical analyses. Q.Q. and K.P performed snDrop-seq and data preprocessing. K.C. performed and analyzed electrophysiology experiments. A.J.V. performed immunohistochemistry experiments. S.D.M., C.N.Q., and K.T.L. scored behavioral tests. M.V.F. led electrophysiology experiments. H.W. led snDrop-seq. Y.D. led chromatin biochemistry work. E.K. led the project.

## STAR METHODS

### RESOURCE AVAILABILITY

#### Lead contact

Further information and requests for resources and reagents should be directed to and will be fulfilled by the lead contact, Dr. Erica Korb.

#### Materials availability

- This study did not generate new unique reagents.

#### Data and code availability

- CUT&Tag, ATAC-seq, RNA-seq, single-nucleus Drop-seq data have been deposited at GEO and are publicly available as of the date of publication. Accession numbers are listed in the key resources table. Original western blot images have been deposited at Mendeley and are publicly available as of the date of publication. The DOI is listed in the key resources table. Microscopy data reported in this paper will be shared by the lead contact upon request.
- This paper does not report any original code.
- Any additional information required to reanalyze the data reported in this paper is available from the lead contact upon request.

## EXPERIMENTAL MODEL AND STUDY PARTICIPANT DETAILS

### Primary neuronal culture

Cortices were dissected from E16.5 C57BL/6J embryos and cultured in supplemented neurobasal medium (Neurobasal [Gibco 21103-049], B27 [Gibco 17504044], GlutaMAX [Gibco 35050-061], Pen-Strep [Gibco 15140-122]) in TC-treated 12- or 6well plates coated with 0.05 mg/mL Poly-D-lysine (Sigma-Aldrich A-003-E). At 3-4 DIV, neurons were treated with 0.5 µM AraC. For all experiments using cultured cortical neurons, neurons were collected at 12 DIV.

### Mice

An H2BE-KO mouse was generated as described previously^18^. In brief, the endogenous H2BE CDS was replaced with a membrane-targeted mCherry reporter sequence in C57Bl/6 mice. All mice were housed in a 12-hour light-dark cycle and fed a standard diet. All experiments were conducted in accordance with and approval of the IACUC. For all experiments, samples were collected from mice between 3-5-months-old and both male and female mice were included, except for single-nucleus Drop-seq. Drop-seq was performed on cortical tissue from male mice only. This mouse is available from The Jackson Laboratory (strain # 023819).

## METHOD DETAILS

### Histone extraction

Snap-frozen brain tissues were homogenized in nuclear isolation buffer (NIB; 15mM Tris-HCl pH 7.5, 60mM KCl, 1mM CaCl_2_, 15mM NaCl, 5mM MgCl_2_, 250mM sucrose supplemented by protease inhibitor [Roche 04693124001], phosphatase inhibitor [Roche 04906837001], 1mM DTT, 1mM PMSF) + 0.3% NP-40 using a pre-chilled dounce and pestle. Sample were then incubated on ice for 5 minutes, and centrifuged for 5 minutes at 1000g at 4°C. The pellet was washed 1x in NIB without NP-40 and centrifuged for 5 minutes at 1000g at 4°C. The pellet was then resuspended in 1mL cold 0.4N H_2_SO_4_ and rotated overnight at 4C. Following the overnight incubation, samples were pelleted for 5 minutes at 10,000g at 4°C and the supernatant was transferred to a new tube. Trichloroacetic acid was added to 25% by volume, and the cells were left on ice at 4°C overnight. Cells were again pelleted for 5 minutes at 10,000g at 4°C, and the supernatant was discarded. Samples were centrifuged for 5 minutes at 16,000g at 4°C and supernatant was discarded. The pellet was washed 2X with ice-cold acetone. After the second wash, samples were air-dried. The pellet was resuspended in molecular biology-grade H_2_O, sonicated in a Biorupter for 10 min, and then incubated at 50°C in a thermomixer shaking at 1000rpm. Samples were centrifuged for 10 minutes at 16,000g at 4°C and the supernatant containing the histone fraction was collected.

### Western blotting

Protein lysates or histone samples were mixed with 5X Loading Buffer (5% SDS, 0.3M Tris pH 6.8, 1.1mM Bromophenol blue, 37.5% glycerol), boiled for 10 minutes, and cooled on ice. Protein was resolved by 16% Tris-glycine SDS-PAGE, followed by transfer to a 0.45-μm PVDF membrane (Sigma-Aldrich IPVH00010) for immunoblotting. Membranes were blocked for 1 hour at RT in 5% milk in 0.1% TBST and probed with primary antibody overnight at 4C. Membranes were incubated with secondary antibody for 1 hour at RT.

### Immunohistochemistry

#### Brain sectioning

Mice were anesthetized with isofluorane and transcardially perfused with 4% paraformaldehyde. Brains were collected, rinsed with ice-cold PBS, placed in 15% sucrose at 4°C and allow to settle to the bottom. Brains were then transferred to 30% sucrose and again allowed to settle to the bottom. Brains were then cryopreserved in O.C.T. Brains were coronally sectioned at 18μm using a cryostat and mounted on Superfrost Plus slides.

#### Staining

Sections were washed once with 1X PBS, then blocked for 1 hour in blocking buffer (10% donkey serum supplemented with 0.5% Triton X-100 in 1X PBS). Sections were washed once with 1X PBS, then incubated with primary antibody (anti-H2BE, 1:500, Sigma ABE1384, anti-NeuN Alexa Fluor 488 conjugated, 1:200, Sigma MAB377X) in a 1:10 dilution of blocking buffer overnight at 4°C in a humidified chamber. Sections were washed 3 times in 1X PBS, incubating for 5 minutes, then incubated with secondary antibody (Donkey anti-rabbit IgG Alexa Fluor 568, 1:500, Thermo A10042) in a 1:10 dilution of blocking buffer in a dark chamber for 1 hour at RT. After three additional washes for 5 minutes each, sections were incubated with 1ug/uL DAPI (Thermo Scientific 62247) for 10 minutes at RT. Coverslips were mounted onto slides using Prolong Gold mounting medium. Slides were imaged on an upright Leica DM 6000, TCS SP8 laser scanning confocal microscope. The objective used was a ×63 HC PL APO CS2 oil objective with an NA of 1.40.

### Quantitative mass spectrometry

#### In-Solution Digestion

Samples were solubilized and digested per the S-Trap Micro (Protifi) manufacturer’s protocol^52^. Briefly, samples were solubilized in 50 µL of extraction buffer containing 5% sodium dodecyl sulfate (SDS, Affymetrix), 50mM TEAB (pH 8.5, Sigma), and protease inhibitor cocktail (Roche cOmplete, EDTA free), reduced in 5mM TCEP (Thermo), alkylated in 20mM iodoacetamide (Sigma), then acidified with phosphoric acid (Aldrich) to a final concentration of 1.2%. Samples were diluted with 90% methanol (Fisher) in 100 mM TEAB, then loaded onto an S-trap column and washed three times with 50/50 chloroform/methanol (Fisher) followed by three washes of 90% methanol in 100 mM TEAB. A 1:10 ratio (enzyme: protein) of Trypsin (Promega) and LysC (Wako) suspended in 20µL 50mM TEAB was added, and samples were digested for 1.5 hours at 47 °C in a humidity chamber. After incubation, peptides were eluted with an additional 40 μL of 50 mM TEAB, followed by 40 μL of 0.1% trifluoroacetic acid (TFA) (Pierce) in water, and finally 40 μL of 50/50 acetonitrile:water (Fisher) in 0.1% TFA. Eluates were combined and desalted directly using Phoenix peptide cleanup kit (PreOmics) per manufactures protocol, dried by vacuum centrifugation and reconstituted in 0.1% TFA containing iRT peptides (Biognosys, Schlieren, Switzerland).

#### Target Assay Development

Parallel Reaction Monitoring (PRM) assay^53,54^ was developed using target peptides identified from data dependent acquisition (DDA) analysis of recombinant murine histone H2BE- and a tryptic digest of enriched murine histones. A peptide unique to H2BE (EIQTSVR) and a peptide common to other H2B variants (EIQTAVR) were selected. Heavy isotope-labelled peptides were synthesized, quantified by amino acid analysis, aliquoted, and lyophilized by JPT Peptide Technologies, Berlin. To test and optimize the PRM method, 8ng of heavy isotope peptides were initially spiked into 250ng of tryptic E.coli digest and injected on column. After PRM method optimization, heavy labelled peptides were spiked into 2ug Histone tryptic so that 5ng was injected on column for each peptide.

#### Mass Spectrometry Data Acquisition

Samples were analyzed on a Q-Exactive HF mass spectrometer (Thermofisher Scientific San Jose, CA) coupled with an Ultimate 3000 nano UPLC system and an EasySpray source. Peptides were loaded onto an Acclaim PepMap 100 75um x 2cm trap column (Thermo) at 5uL/min and separated by reverse phase (RP)-HPLC on a nanocapillary column, 75 μm id × 50cm 2um PepMap RSLC C18 column (Thermo). Mobile phase A consisted of 0.1% formic acid and mobile phase B of 0.1% formic acid/acetonitrile. Peptides were eluted into the mass spectrometer at 300 nL/min with each RP-LC run comprising a 90-minute gradient from 3% B to 45% B.

The mass spectrometer was set to repetitively scan m/z from 300 to 1400 (R = 240,000) followed by data-dependent MS/MS scans on the twenty most abundant ions, minimum AGC 1e4, dynamic exclusion with a repeat count of 1, repeat duration of 30s, and resolution of 15000. The AGC target value was 3e6 and 1e5, for full and MSn scans, respectively. MSn injection time was 160 ms. Rejection of unassigned and 1+,6-8 charge states was set.

The parallel reaction monitoring (PRM) method^53,54^ combined two scan events. For the full scan we used 150– 2000 m/z mass selection, resolution 120,000 at m/z 200, automatic gain control (AGC) target value of 3e6, and maximum injection time of 200ms. The targeted MS/MS was run at an Orbitrap resolution of 30,000 at m/z 200, an AGC target value of 5e6, and maximum fill time of 200ms. and a target isolation window of 1.2 m/z. Fragmentation was performed with normalized collision energy (NCE) of 27 eV. Table 1 shows targeted inclusion list with scheduled retention times for heavy and light versions of EIQTAVR and EIQTSVR., along with unscheduled acquisition of iRT peptides for internal quality control.

**Table 1.** The PRM scheduled list for heavy and light peptides.

| Mass<br>[m/z] | Formula<br>[M] | Species [z] | Polarity | Start[min] | End[min] | N(CE) | MSX | ID | Comment |
| --- | --- | --- | --- | --- | --- | --- | --- | --- | --- |
| 408.73233 |  | 2 | Positive | 25 | 35 | 27 |  |  | EIQTAVR(light) |
| 413.73647 |  | 2 | Positive | 25 | 35 | 27 |  |  | EIQTAVR(heavy) |
| 416.72979 |  | 2 | Positive | 25 | 35 | 27 |  |  | EIQTSVR(light) |
| 421.73393 |  | 2 | Positive | 25 | 35 | 27 |  |  | EIQTSVR(heavy) |

**Table 2.** Marker genes used to determine cell identity in Drop-seq.

| Cell identity | Marker gene(s) | References |
| --- | --- | --- |
| Endothelial cells | Flt1 | 29-30,32 |
| Microglia | Ctss | 29-30,32 |
| Astrocytes | Gja1 | 29-30,32 |
| Oligodendrocytes | Mog | 29,32 |
|  | Enpp6 | 29-30,32 |
|  | Opalin | 30,32 |
| Excitatory neurons | Slc17a7 | 29-32 |
| Inhibitory Neurons | Gad2 | 29-31 |
| Cortical layer 2/3 neurons | Ndst4 | 30 |
|  | Enpp2 | 29-30 |
| Cortical layer 4 neurons | Rorb | 29-33 |
| Cortical layer 5 neurons | Tox | 30,33 |
| Cortical layer 6 neurons | Zfpm2 | 30 |
|  | Hs3st4 | 30 |
| Cortical layer 6b neurons | Ctgf | 30,33 |
| Clastrum | Gnb4 | 29-30 |

#### System Suitability and Quality Control

The suitability of Q Exactive HF instrument was monitored using QuiC software (Biognosys, Schlieren, Switzerland) and Skyline^55^ for the analysis of the spiked-in iRT peptides in each sample. Meanwhile, as a measure for quality control, we injected standard *E. coli* protein digest prior to and after injecting sample set and collected the data in the Data Dependent Acquisition (DDA) mode. The collected data were analyzed in MaxQuant^56^ and the output was subsequently visualized using the PTXQC^57^ package to track the quality of the instrumentation.

#### MS data processing and analysis

The raw files for DDA analysis were processed with MaxQuant version 2.0.1.0 using its default workflow. The reference *E. coli* proteome from UniProt was concatenated with murine histones and common protein contaminants and used for the raw data search.

Skyline 21.2.0.568 with its default settings was used to process the PRM raw data. The heavy-isotope label for the C-terminal Arg was set as static modification. All peak integrations were reviewed manually and the sum of all transitions was calculated for light and heavy peptides. For each sample, light/heavy peak area ratio was calculated and multiplied by the known heavy peptide amount per injection to estimate the light peptide abundance value.

### CUT&Tag-sequencing

#### Library preparation & sequencing

Input samples were 400K primary cortical neurons per biological replicate. CUT&Tag was performed according to published protocols^58^. Prior to sequencing, library size distribution was confirmed by capillary electrophoresis using an Agilent 4200 TapeStation with high sensitivity D1000 reagents (5067-5585), and libraries were quantified by qPCR using a KAPA Library Quantification Kit (Roche 07960140001). Libraries were sequenced on an Illumina NextSeq550 instrument (42-bp read length, paired end).

#### Data processing and analysis

Reads were mapped to *Mus musculus* genome build mm10 with Bowtie 2 (v2.4.5)^59^. Six million reads from each biological replicate were subset and each condition was then merged across biological replicates (SAMtools^60^ v1.15). Heatmaps were generated using deepTools^61^ (v3.5.1). Metaplots were generated using ngs.plot^62^ (v2.63) against the mouse genome. Peaks were called using MACS3^63^ (v3.0.0b1) and annotated using Homer^64^ (v4.10) . For downstream analysis, we used a Peak Score cutoff of 25 and removed peaks that were assigned to ‘ChrUn’ (unknown chromosome) by Homer. The R package GenomicDistributions^65^ (v1.6.0) was used to analyze the genomic distribution of peaks. IGV tools^66^ (2.12.3) was used to generate genome browser views.

To compare CUT&Tag signal to gene expression, normalized read counts from RNA-sequencing of WT primary neuronal cultures (see RNA-sequencing methods section) were used to generate gene lists by expression level. Genes with base mean < 3 were defined as ‘not expressed’. The remaining genes comprise the ‘all expressed’ categorization, and these genes were further divided into 3-quantiles (by base mean) to define ‘low expression’, ‘mid expression’ and ‘high expression’.

#### Gene ontology

For gene ontology analysis, gene names were assigned to peak coordinates using Homer. Peaks that were annotated as ‘intergenic’, ’non-coding’, and ‘NA’ by Homer were not included in GO analysis. PANTHER^67,68^ (v18.0) was used to perform an overrepresentation test against the biological process complete ontology using default parameters. The *Mus musculus* genome was used as a background gene list. For conciseness and visualization, parent terms were excluded and only the most specific GO terms were plotted.

### ChIP-sequencing

#### Chromatin immunoprecipitation

After 12 DIV, cells were fixed with 1% PFA in PBS for 7min and the reaction was quenched with 2.5M glycine. Cells were washed twice with PBS, collected in PBS (with protease inhibitor [Roche 04693124001], phosphatase inhibitor [Roche 04906837001], 1mM DTT, 1mM PMSF), and then pelleted at 1,200 rpm for 5lllmin. Cells were then rotated in lysis buffer 1 (50lllmM HEPES-KOH pHlll7.5, 140lllmM NaCl, 1lllmM EDTA, 10% glycerol, 0.5% NP-40 and 0.25% Triton X-100) for 10lllmin at 4°C and spun at 1,350 g for 5lllmin at 4lll°C to isolate nuclei. Supernatant was discarded and cells were resuspended in lysis buffer 2 (10lllmM Tris-HCl pHlll8, 200lllmM NaCl, 1lllmM EDTA and 0.5lllmM EGTA) to lyse nuclei. Samples were rotated for 10lllmin at room temperature and were spun again at 1,350 g for 5lllmin at 4°C. The supernatant was discarded, and the pellet was resuspended in lysis buffer 3 (10lllmM Tris-HCl pHlll8, 100lllmM NaCl, 1lllmM EDTA, 0.5lllmM EGTA, 0.1% sodium deoxycholate and 0.5% N-lauroylsarcosine). Lysates were sonicated using a Covaris S220 Focused-ultrasonicator for 45lllmin (peak power: 140; duty factor: 5.0; 200 cycles per burst; avg power: 7.0). Triton X-100 was added to reach a final concentration of 1%, and lysates were spun at 18,000 g for 10lllmin at 4°C. 50lllµl of lysate was saved as input shearing control.

Antibody-conjugated Protein A Dynabeads (2.5µg of antibody conjugated to 75lllµl of Protein A Dynabeads, resuspended in 50lllµl per immunoprecipitation) were added to the lysates overnight with rotation at 4°C. Beads were then washed 8X with RIPA wash buffer (50lllmM HEPES-KOH pHlll7.5, 500lllmM LiCl, 1lllmM EDTA, 1% NP-40 and 0.7% sodium deoxycholate) and 1X with TE + 50lllmM NaCl. Chromatin was eluted from beads for 45lllmin with shaking at 65°C in elution buffer (50lllmM Tris-HCl pHlll8.0, 10lllmM EDTA and 1% SDS). Samples were removed from beads and cross-linking was reversed in both input and IP samples by incubating overnight at 65lll°C. RNA was digested with 5ug/mL RNase for 45 min at 37lll°C, and protein was digested with 0.2ug/mL proteinase K for 1 hr at 55lll°C. DNA was then purified with the Zymo DNA Clean & Concentrator Kit with a final elution in 50uL molecular biology grade H2O.

#### Library preparation & sequencing

Sequencing libraries were prepared using the TruSeq ChIP Library Preparation Kit (Illumina 15023092). Prior to sequencing, library size distribution was confirmed by capillary electrophoresis using an Agilent 4200 TapeStation with high sensitivity D1000 reagents (5067-5585), and libraries were quantified by qPCR using a KAPA Library Quantification Kit (Roche 07960140001). Libraries were sequenced on an Illumina NextSeq1000 instrument (66-bp read length, paired end).

#### Data processing and analysis

Reads were mapped to *Mus musculus* genome build mm10 with Bowtie 2 (v2.4.5)^59^. Thirty million reads from each biological replicate were subset and each condition was then merged across biological replicates (SAMtools^60^ v1.15). Metaplots were generated using ngs.plot^62^ (v2.63) against the mouse genome. The R package GenomicDistributions^65^ (v1.6.0) was used to analyze the genomic distribution of peaks.

### Dot Blot

0.2-μm PVDF paper was immersed in methanol and placed on TBS-soaked filter paper. Peptides were dotted onto the PVDF paper at serial dilutions and allowed to dry for 4 hours. The membrane was then wet in methanol again and equilibrated in TBS before blocking for 1 hour in 3% non-fat dry milk dissolved in TBS. The membrane was then incubated in antibody solution for 1 hour at room temperature and washed 3 times for five minutes in TBS-T (TBS with 0.1% Tween-20) before incubation with secondary antibody for 1 hour. Following secondary, the membrane was again washed 3 times for five minutes before imaging.

### Protein extraction

Tissues were homogenized in 500uL RIPA buffer (10% sucrose, 1% SDS, 5mM HEPES pH 7.9, 10mM sodium butyrate in MilliQ water supplemented by protease inhibitor [Roche 04693124001], phosphatase inhibitor [Roche 04906837001], 1mM DTT, 1mM PMSF) using a pre-chilled dounce and pestle. Samples were then titrated 5x with a 26.3 gauge needle and centrifuged for 15 min at max speed at 4°C. Supernatant containing protein lysates were transferred to a new tube.

### Chromatin Biochemistry Methods

#### DNA preparation for chromatin assembly

Mononucleosome-sized 601 DNA was prepared by PCR amplification of a DNA template containing one copy of the Widom 601 DNA sequence^69^. PCR products were then pooled and purified with a QIAquick PCR Purification Kit (Qiagen), using water to elute from the final columns. Eluents were pooled, frozen, and lyophilized before being resuspended in buffer TE (10 mM Tris-HCl pH 7.6, 0.1 mM EDTA), quantified by NanoDrop OneC, and adjusted to a final concentration of approximately 1-1.5 g/L.

DNA templates used for chromatin fibers were prepared essentially as described before^70^. *E. coli* DH5α cells were transformed with a pWM530 vector bearing 12 repeats of the Widom 601 nucleosome positioning sequence with a 30 bp linker and used to inoculate 6 L of luria broth under ampicillin selection and grown at 37 °C for 18-24 hrs. Cultures were harvested and DNA purified using a Plasmid Giga Kit (Qiagen). Purified DNA was resuspended in buffer TE and adjusted to a concentration of ∼1 g/L. Plasmid was then digested with EcoRI and EcoRV (NEB) overnight to generate linear chromatin templates (i.e., dslDNA or 601^18^-L. Complete digestion was determined by agarose gel electrophoresis in 1X TAE buffer. Next, digestion reactions were fractionated by PEG 6000-induced precipitation as described previously to separate the ∼3.2 kb desired chromatin substrates from the 200-300 bp plasmid backbone fragments^71^. Precipitated DNA was next resuspended in buffer TE, purified by phenol/chloroform extraction, and precipitated with absolute ethanol as described^72^. DNA was finally resuspended in buffer TE, quantified by NanoDrop OneC, and adjusted to a concentration of 1-1.5 g/L. Buffer DNA was likewise prepared by digestion of a similar construct containing 8 repeats of the 155 bp mouse mammary tumor virus (MMTV) weak nucleosome positioning sequence with EcoRV, PEG precipitation, and phenol/chloroform extraction.

#### Histone expression and purification

Recombinant human histones were expressed and purified as previously described^69^. Briefly, H2A, H2B, H2BE, H3.1, H4, and histone mutants H2B V39I and H2BE I39V were expressed in *E. coli* BL21(DE3) or C41(DE3)) cells at 37 °C until reaching an OD_600_ of 0.6-0.8, before induction with 500 µM IPTG for 4 hours at 37 °C. Bacteria were then harvested by centrifugation at 6000 x g for 25 minutes.

Cell pellets were resuspended in lysis buffer (200 mM NaCl, 20 mM Tris-HCl pH 7.6, 1 mM EDTA, 1 mM β-mercaptoethanol) and then lysed via sonication. Lysate was cleared by centrifugation at 30,000 x g for 20 minutes and the remaining insoluble pellet was resolubilized in extraction buffer (6 M guanidine HCl, 1 mM DTT, 1x PBS, pH 7) and nutated at 4 °C overnight. The extraction was cleared by centrifugation at 30,000 x g for 40 minutes and filtered using a 0.45-micron syringe filter (Fisher Scientific). Filtered extraction was diluted 1:1 with HPLC buffer A (0.1% trifluoroacetic acid (TFA) in water) before purification with reverse-phase HPLC on a 30-70% buffer B (0.1% TFA in 90% acetonitrile and 10% water). Absorbance at 280 nm was used to observe desired peaks and fractions were collected using an automated fraction collector. Purity of fractions was determined using a quadrupole LC-MS. Fractions deemed pure were combined, lyophilized and stored at - 80 °C.

#### Histone octamer assembly

To assemble histone octamers, monomeric core histones were resuspended in unfolding buffer (20 mM Tris-HCl pH 7.6, 6 M guanidine hydrochloride, 1 mM DTT) and quantified by measuring A280 using a NanoDrop. Core histones were next combined at the following stoichiometries to generate a slight excess of H2A/H2B dimer to assist with subsequent purification: 1.05:1.05:1:1 H2A:H2B:H3:H4. Total protein concentration of the mixture was adjusted to 1 g/L, and samples were dialyzed against octamer refolding buffer (10 mM Tris-HCl pH 7.6, 2 M NaCl, 1 mM EDTA, 1 mM DTT) at 4 °C using 3.5K MWCO Slide-A-Lyzer dialysis cassettes. A total of three rounds of dialysis against refolding buffer were performed, with the first exchange going overnight and the subsequent two lasting at least 6 hrs.

Following the final dialysis, samples were harvested from dialysis cassettes, cleared by centrifugation, and concentrated using 30K MWCO Amicon centrifugal filter units. Finally, samples were centrifuged at 17,0000 xg for a minimum of 10 minutes at 4 °C before being injected onto an AKTA 25L FPLC instrument and resolved over a SuperDex 200 10/300 Increase column, using octamer refolding buffer for the liquid phase. Fractions were analyzed by SDS-PAGE on 12 % acrylamide gels, and those containing octamers were pooled and concentrated using Amicon 30K MWCO centrifugal filter unites. Octamers were quantified by A280, adjusted to 50 % (v/v) glycerol, and stored at -20 °C for future use.

#### Chromatin reconstitution

Nucleosome core particles (NCPs), linear nucleosome arrays, and circular nucleosome arrays were all prepared by the same method of salt gradient dialysis with all steps at 4 °C. Substrate DNA and recombinant histone octamers were combined in approximately equimolar quantities (with respect to the expected nucleosome load of the DNA, i.e., 1 for mononucleosomal DNA, or 12 for chromatin arrays) with final buffer conditions identical to octamer refolding buffer – 10 mM Tris-HCl pH 7.6, 2 M NaCl, 0.1 mM EDTA, 1 mM DTT. For nucleosome array assemblies MMTV DNA was also added, but at a lower stoichiometry to act as a buffer that facilitates array assembly without competing for octamer occupancy (0.2:1 MMTV:NCP). Optimal assembly stoichiometries were determined empirically for all templates, with Octamer:DNA stoichiometries being approximately 1.2:1 for mononucleosomes and 1.6:1 for 601 nucleosome arrays.

Assembly reactions were combined and mixed by gentle pipetting, centrifuged for 5 min at 17,000 xg at 4 °C, and then added to Slide-A-Lyzer Mini dialysis buttons pre-moistened in array initial buffer (10 mM Tris-HCl pH7.6, 1.4 M NaCl, 0.1 mM EDTA, 1 mM DTT) and dialyzed against 200 mL of the same buffer for 1 hr. Next, a peristaltic pump was used to transfer 350 mL of dilution buffer (10 mM Tris-HCl pH 7.6, 10 mM NaCl, 0.1 mM EDTA, 1 mM DTT) to the samples in initial buffer with a flow rate of 0.5-1.0 mL/min. After all dilution buffer was transferred, arrays were left to dialyze for at least 1 hr or up to overnight. Samples were next moved to a fresh 350 mL of dilution buffer and dialyzed for 6 hrs. Lastly, samples were moved 300 mL of fresh dilution buffer and dialyzed for 1-2 hrs before harvesting.

Mononucleosome samples were simply pipetted out of dialysis buttons, centrifuged for 5 min at 17,000 xg, and supernatant (in case any precipitation was present) was quantified, subject to EMSA to validate assembly efficiency, and stored at 4 °C for up to 2 months. Chromatin array samples were similarly harvested to yield a mixture of assembled chromatin fibers, MMTV mononucleosomes, and free MMTV DNA. To separate the chromatin fibers, an equal volume of precipitation buffer (10 mM Tris-HCl pH 7.6, 10 mM NaCl, 10 mM MgCl_2_) was added to samples followed by a 20-minute incubation on ice. Next, samples were centrifuged for 10 minutes at 17,000 xg. Supernatant was gently pipetted off samples to avoid disturbing the barely visible pellets, to which a desired volume of array dilution buffer was next added. Pellets were left on ice for 10 minutes undisturbed to allow for the gradual resuspension of chromatin pellets before being quantified.

Finally, EMSA was used to validate assemblies and harvests. Chromatin arrays were stored at 4 °C for up to one week before use.

#### Electrophoretic mobility shift assays (EMSAs)

Mononucleosome EMSAs were performed using polyacrylamide gel electrophoresis in 5 % acrylamide, 0.5 X TBE gels. A solution of 1 M sucrose was used as a loading buffer (1:3 dilution) for nucleosome samples. Gels were run for 30-40 minutes at a constant voltage of 130 V, stained with SYBR Gold dye diluted 1:10,000 in 0.5 X TBE buffer for 5-10 minutes, and DNA migration was visualized on an Amersham AI600 imager (GE/Cytiva) using the UV 312 nm channel. If needed, protein migration was visualized after DNA imaging by staining gels with Imperial Protein Stain and visualized on the AI600 instrument in the colorimetric channel.

Chromatin array EMSAs were performed similarly, except with a different gel formulation. Agarose-polyacrylamide gel electrophoresis (APAGE) gels were cast using the Mini-PROTEAN Tetra Handcast system (Bio-Rad). Before combining gel-casting reagents, the gel-facing sides of the 1.5 mm spacer plates, short plates, and 10-well gel combs were lubricated with a thin layer of 50 % (v/v) glycerol. Next, ultrapure water, 50X TAE (sufficient for a final concentration of 0.5 X), and powdered agarose (sufficient for a final concentration of 1 % w/v) were heated until dissolved. This solution was rapidly combined with 40 % acrylamide (37.5:1 mono:bis) solution (Bio-Rad) (sufficient for a final concentration of 2 %), APS (sufficient for 0.125 %), and TEMED (sufficient for 0.04 %), and added into the assembled gel casting cassettes. Gels were allowed to cool and polymerize for at least 1 hr. Once cool, gels were pre-run at 4 °C in 0.5 X TAE buffer for 3 hrs at 100 V. Next, chromatin samples could be loaded onto gels using sucrose loading solution and the gels run for approximately 1 hr at 120 V at room temperature before visualization as mononucleosome EMSAs.

#### Atomic force microscopy

DNA-protein complexes were imaged using an Asylum MFP 3D Bio AFM (Oxford Instruments, Goleta CA) with an Olympus AC240TS probe in tapping mode at room temperature. The samples were prepared a suitable concentration (0.5-1.0 ng/µL), then 40 uL of prepared samples were slowly deposited to a freshly cleaved AP-mica for 5 minutes and rinsed with 1 mL ultrapure deionized water twice before being gently dried with UHP argon gas.

AFM images were collected at a speed of 0.5-1 Hz at 512 × 512-pixel resolution, with an image size of 2 μm. For analysis, raw images were exported into 8-bit grayscale Tiff images using the Asylum Research’s Igor Pro software and imported into FIJI/ImageJ (NIH) for detection of single particles and quantification of volume, surface area, and volume/surface area ratio using as has been done previously for studies of chromatin compaction via AFM^25^. In order to assess single chromatin particles, rather than potential clusters of multiple fibers or residual MMTV mononucleosomes, only particles with volumes measuring between 2,000 and 15,000 nm^3^ were included in analyses.

#### Magnesium-dependent self-association assay

Chromatin compaction was tested by magnesium-drive self-association as described previously^26^. Briefly, a magnesium solution (100 mM MgCl_2_, 10 mM Tris-HCl pH 7.6, 10 mM NaCl) was titrated into the sample solution to raise the magnesium concentration in 0.5 mM increments. After each addition, samples were allowed to sit on ice for 10 min and then centrifuged for an additional 10 min at 17,000 xg at 4 °C. The concentration of soluble DNA from nucleosome arrays was measured by NanoDrop.

#### Nucleosome differential scanning fluorimetry (DSF) assays

Nucleosome stability was measured using a Protein Thermal Shift kit (Applied Biosystems) with the following modifications. The assay was conducted at 10 μL volumes with 10X SYPRO Orange dye (Invitrogen) and nucleosome dilution buffer (10 mM Tris-HCl pH 7.6, 100 mM NaCl, 0.1 mM EDTA, and 1 mM DTT) in 384-well plates. Final nucleosome concentrations used were approximately 60 ng/μL of DNA. Fluorescence melt curve data was acquired using a QuantStudio 5 Real-Time PCR System (Applied Biosystems) with the following method: initial ramp rate of 1.6 °C/s to 25 °C with a 5-minute hold time at 25 °C, followed by a second ramp rate of 0.05 °C/s to 99.9 °C with a 2-minute hold time at 99.9 °C. Melting temperatures were calculated using the Protein Thermal Shift software (Applied Biosystems).

### ATAC-sequencing

#### Library preparation & sequencing

Input samples were 400K primary cortical neurons per biological replicate. Neurons were collected by scraping in cold lysis buffer (10lllmM Tris-Cl (pHlll7.5), 10lllmM NaCl, 3lllmM MgCl_2_, 0.1% (vol/vol) NP-40, 0.1% (vol/vol) Tween-20 and 0.01% (vol/vol) digitonin) and washed in wash buffer (10lllmM Tris-Cl (pHlll7.5), 10lllmM NaCl, 3lllmM MgCl_2_ and 0.1% (vol/vol) Tween-20). Transposition was performed with Tagment DNA TDE1 (Illumina, 15027865). Transposition reactions were cleaned with AMPure XP beads (Beckman, A63880), and libraries were generated by PCR with NEBNext High-Fidelity 2× PCR Master Mix (NEB, M0541). Prior to sequencing, library size distribution was confirmed by capillary electrophoresis using an Agilent 4200 TapeStation with high sensitivity D1000 reagents (5067-5585), and libraries were quantified by qPCR using a KAPA Library Quantification Kit (Roche 07960140001). Libraries were sequenced on an Illumina NextSeq550 instrument (42-bp read length, paired end).

#### Data processing and analysis

Reads were mapped to *Mus musculus* genome build mm10 with Bowtie 2 (v2.4.5)^59^. Fifty million reads were subsetted from each biological replicate and each condition was then merged across biological replicates (SAMtools^60^ v1.15). Heatmaps were generated using deepTools^61^ (v3.5.1). Metaplots were generated using ngs.plot^62^ (v2.63) against the mouse genome. Peaks were called using MACS3^63^ (v3.0.0b1) and annotated using Homer^64^ (v4.10) . For downstream analysis, we used a Peak Score cutoff of 25 and removed peaks that were assigned to ‘ChrUn’ (unknown chromosome) by Homer. Differential ATAC peaks were called using DiffBind^73^ (v3.4.11). The following genomic regions were used as input data for DiffBind analysis: WT ATAC peak coordinates from MACS3 (Fig 2D-G), H2BE binding site coordinates from MACS3 peak calling of CUT&Tag data (Fig 2H-K), TSS coordinates +/-500bp from mm10 (Fig S2D-F), enhancer regions from brain tissue (Fig S2G-I)^74^. IGV tools^66^ (2.12.3) was used to generate genome browser views.

To compare ATAC signal within different genomic regions, we subset H2BE binding sites based on their Homer annotations into promoter (‘promoter-TSS’ in Homer), gene body (‘exon’ and ‘intron’ in Homer), and intergenic (‘intergenic’ in Homer).

### Viral infection

#### Viral constructs

The GFP control plasmid was obtained from Addgene, pLenti-CMV-MCS-GFP-SV-puro (Addgene plasmid 73582).

The H2BE and H2B constructs were generated as previously described^18^ and were provided by the Dulac lab. The H2B and H2BE coding sequences were then moved to the pLenti backbone through Gibson assembly. Primers were designed through NEB Builder to separately amplify H2B or H2BE and the pLenti backbone. Each primer contained a non-complementary region on its 5’-end that corresponded to approximately 10 bases on the other template. PCR amplification was performed with Q5 High-Fidelity DNA Polymerase (NEB M0491S). Following DPN1 digestion, fragments were ligated together using NEBuilder HiFi DNA Assembly Master Mix (NEB E2621S) and transformed into NEB 5-alpha Competent *E. coli* cells (E2621S). Plasmid sequence was verified through Plasmidsaurus long read sequencing.

The H2BE-I39V and H2BE-L3P constructs were generated by site-directed mutagenesis of the H2BE backbone using Pfu Turbo HotStart DNA polymerase (Agilent, 600322-51), and primers were created using the DNA-based primer design feature of the online PrimerX tool. Constructs were verified by Sanger sequencing.

#### Lentiviral production

HEK293T cells were cultured in high-glucose DMEM growth medium (Corning 10-013-CV), 10% FBS (Sigma-Aldrich F2442-500ML), and 1% Pen-Strep (Gibco 15140-122). Calcium phosphate transfection was performed with Pax2 and VSVG packaging plasmids. Viral media was removed 2 h after transfection and collected at 48 and 72 h later. Viral media was passed through a 0.45-μm filter and precipitated for 48 hours with PEG-it solution (40% PEG-8000 [Sigma-Aldrich P2139-1KG], 1.2 M NaCl [Fisher Chemical S271-1]). Viral particles were pelleted and resuspended in 200μL PBS.

#### Neuronal infection

At 10 DIV, neurons were transduced overnight with lentivirus containing the constructs described above. Virus was removed the following day, and neurons were cultured for one additional day.

### RNA-sequencing

#### Library preparation & sequencing

Input samples were 400 primary cortical neurons from 4 WT biological replicates and 4 KO biological replicates. RNA was isolated using Zymo Quick-RNA Miniprep Plus Kit (R1057). Prior to library preparation, RNA integrity was confirmed using an Agilent 4200 TapeStation with high sensitivity RNA reagents (5067-5579). Sequencing libraries were prepared using the TruSeq Stranded mRNA kit (Illumina 20020595). Prior to sequencing, library size distribution was confirmed by capillary electrophoresis using an Agilent 4200 TapeStation with high sensitivity D1000 reagents (5067-5585), and libraries were quantified by qPCR using a

KAPA Library Quantification Kit (Roche 07960140001). Libraries were sequenced on an Illumina NextSeq1000 instrument (66-bp read length, paired end).

#### Data processing and analysis

Reads were mapped to *Mus musculus* genome build mm10 with Star^75^ (v2.7.9a). The R packages DESeq2^76^ (v1.38.3) and limma (v3.54.2) via edgeR^77^ (v3.40.2) were used to perform differential gene expression analysis. We defined genes as differentially expressed where FDR < 0.05 and absolute fold change >= 1.25. Volcano plots were generated using VolcaNoseR^78^. IGV tools^66^ (2.12.3) was used to generate genome browser views.

#### Gene ontology

PANTHER^67,68^ (v18.0) was used to perform an overrepresentation test against the biological process complete ontology using default parameters. SynGO^28^ was used for synaptic gene ontologies and overrepresentation tests of differentially expressed genes. All expressed genes (defined as base mean 3) was used as a background gene list. For conciseness and visualization, parent terms were excluded and only the most specific GO terms were plotted.

### Single-nucleus Drop-sequencing (sNucDrop-seq)

#### Nuclei isolation

Snap-frozen brain tissues were homogenized in 1mL Buffer A (0.25M sucrose, 50mM Tris-HCl pH7.4, 25mM KCl, 5mM MgCl_2_ supplemented by EDTA-free protease inhibitor [Roche 4693159001]) using a pre-chilled dounce and pestle. Homogenate was then transferred to a pre-chilled 15mL conical tube and mixed with 6mL Buffer B (2.3M sucrose, 50mM Tris-HCl pH7.4, 25mM KCl, 5mM MgCl_2_). An additional 2mL Buffer A was used to rinse leftover homogenate from the dounce and combined with the sample. The homogenate was gently transferred to a pre-chilled 15mL ultracentrifuge tube containing 2mL Buffer C (1.8M sucrose, 50mM Tris-HCl pH7.4, 25mM KCl, 5mM MgCl_2_). Nuclei were pelleted at 100,000 x g for 1.5hr at 4C using a SWI41 rotor. The supernatant was discarded and 1.5mL Buffer D (0.01% BSA in 1X PBS with 0.5U/uL RNase inhibitor [Lucigen 30281-2]) was gently added to the nuclei pellet and incubated on ice 20min. The nuclei pellet were resuspended and the suspension was transferred to a 1.5mL lo-bind tube.

#### Library preparation and sequencing

The single-nucleus suspensions were individually diluted to a concentration of 100 nuclei/mL in DPBS containing 0.01% BSA. Approximately 1.5 mL of this single-nucleus suspension was loaded for each sNucDrop-seq run. The single-nucleus suspension was then co-encapsulated with barcoded beads (ChemGenes) using an Aquapel-coated PDMS microfluidic device (mFluidix) connected to syringe pumps (KD Scientific) via polyethylene tubing with an inner diameter of 0.38mm (Scientific Commodities) (Macosko et al., 2015). Barcoded beads were resuspended in lysis buffer (200 mM Tris-HCl pH8.0, 20 mM EDTA, 6% Ficoll PM-400 (GE Healthcare/Fisher Scientific), 0.2% Sarkosyl (Sigma-Aldrich), and 50 mM DTT (Fermentas; freshly made on the day of run) at a concentration of 120 beads/mL. The flow rates for nuclei and beads were set to 4,000 mL/hr, while QX200 droplet generation oil (Bio-rad) was run at 15,000 mL/hr. A typical run lasts 20 min. Droplet breakage with Perfluoro-1-octanol (Sigma-Aldrich), reverse transcription and exonuclease I treatment were performed, as previously described^29^, with minor modifications. For up to 120,000 beads, 200 μL of reverse transcription (RT) mix (1x Maxima RT buffer (ThermoFisher), 4% Ficoll PM-400, 1 mM dNTPs (Clontech), 1 U/mL RNase inhibitor, 2.5 mM Template Switch Oligo (TSO: AAGCAGTGGTATCAACGCAGAGTGAATrGrGrG), and 10 U/ mL Maxima H Minus Reverse Transcriptase (ThermoFisher)) were added. The RT reaction was incubated at room temperature for 30min, followed by incubation at 42C for 120 min. To determine an optimal number of PCR cycles for amplification of cDNA, an aliquot of 6,000 beads was amplified by PCR in a volume of 50 μL (25 μL of 2x KAPA HiFi hotstart readymix (KAPA biosystems), 0.4 μL of 100 mM TSO-PCR primer (AAGCAGTGGTATCAACGCAGAGT, 24.6 μL of nuclease-free water) with the following thermal cycling parameter (95C for 3 min; 4 cycles of 98C for 20 sec, 65C for 45 sec, 72C for 3 min; 9 cycles of 98C for 20 sec, 67C for 45 sec, 72C for 3 min; 72C for 5 min, hold at 4C). After two rounds of purification with 0.6x SPRISelect beads (Beckman Coulter), amplified cDNA was eluted with 10 μL of water. 10% of amplified cDNA was used to perform real-time PCR analysis (1 μL of purified cDNA, 0.2 μL of 25 mM TSO-PCR primer, 5 μL of 2x KAPA FAST qPCR readymix, and 3.8 μL of water) to determine the additional number of PCR cycles needed for optimal cDNA amplification (Applied Biosystems QuantStudio 7 Flex). We then prepared PCR reactions per total number of barcoded beads collected for each sNucDrop-seq run, using 6,000 beads per 50-μL PCR reaction, and ran the aforementioned program to amplify the cDNA for 4 + 10 to 12 cycles. We then tagmented cDNA using the Nextera XT DNA sample preparation kit (Illumina, FC-131-1096), starting with 550 pg of cDNA pooled in equal amounts, from all PCR reactions for a given run. Following cDNA tagmentation, we further amplified the tagmented cDNA libraries with 12 enrichment PCR cycles using the Illumina Nextera XT i7 primers along with the P5-TSO hybrid primer (AATGATACGGCGACCACCGAGATCTACACGCCTGTCCGCGGAAGCAGTGGTATCAACGCAGAGT*A*C)^79^. After quality control analysis by Qubit 3.0 (Invitrogen) and a Bioanalyzer (Agilent), libraries were sequenced on an Illumina NextSeq 500 instrument using the 75-cycle High Output v2 Kit (Illumina). We loaded the library at 2.0 pM and provided Custom Read1 Primer (GCCTGTCCGCGGAAGCAGTGGTATCAACGCAGAGTAC) at 0.3 mM in position 7 of the reagent cartridge. The sequencing configuration was 20 bp (Read1), 8 bp (Index1), and 60 bp (Read2).

#### Preprocessing of sNucDrop-seq data

Paired-end sequencing reads of sNucDrop-seq were processed using publicly available the Drop-seq Tools v1.12 software^79^ with some modifications. Briefly, each mRNA read (read2) was tagged with the cell barcode (bases 1 to 12 of read 1) and unique molecular identifier (UMI, bases 13 to 20 of read 1), trimmed of sequencing adaptors and poly-A sequences, and aligned using STAR v2.5.2a to the mouse reference genome assembly (mm10, Gencode release vM13). Because a substantial proportion (∼50%) of reads derived from nuclear transcriptomes of mouse cortices were mapped to the intronic regions, the intronic reads were retained for downstream analysis. A custom Perl script was used to retrieve both exonic and intronic reads mapped to predicted strands of annotated genes^29^. Uniquely mapped reads were grouped by cell barcodes. Cell barcodes were corrected for possible bead synthesis errors, using the DetectBeadSynthesisErrors program from the Drop-seq Tools v1.12 software. To generate digital expression matrix, a list of UMIs in each gene (as rows), within each cell (as columns), was assembled, and UMIs within ED = 1 were merged together. The total number of unique UMI sequences was counted, and this number was reported as the number of transcripts of that gene for a given nucleus.

#### Nuclei clustering and marker gene identification

Raw digital expression matrices were combined and loaded into the R package Seurat^80^. For normalization, UMI counts for all nuclei were scaled by library size (total UMI counts), multiplied by 10,000 and transformed to log scale. Nuclei with a relatively high percentage of UMIs mapped to mitochondrial genes (0.2) were discarded. Moreover, nuclei with <=500 UMI or =5000 UMI, discarded, as were nuclei with <=250 genes. Only genes found to be expressing in 10 cells were retained. The Seurat object was then normalized and transformed using the Seurat functions NormalizeData and SCTransform for each genotype. The Seurat functions SelectIntegrationFeatures (nfeatures = 3000), PrepSCTIntegration, FindIntegrationAnchors and IntegrateData were used to integrate the datasets based on the top 3000 more variable features. Prior to clustering, we performed principal component analysis using the RunPCA function and selected 30 principal components for UMAP non-linear dimensional reduction. Based on UMAP, twenty clusters were identified using the Seurat functions Find Neighbors (dim = 30) and FindClusters (resolution = 0.5). To identify marker genes for each cluster, differential expression analysis was performed using the Seurat function FindAllMarkers. Differentially expressed genes that were expressed at least in 25% cells within the cluster and with a fold change more than 0.25 (log scale) were considered marker genes. Cell identity was determined using well-established marker genes for major cortical cell types as described in detail in the table below.

Marker gene analysis led to the identification of 15 cortical neuron clusters (10 excitatory, 5 inhibitory), 1 subcortical neuron cluster, and 4 non-neuronal clusters. Neuronal clusters were annotated according to the cortical layer they occupy, or—if unidentifiable by cortical layer—according to the gene most differentially expressed in that cluster relative to all other excitatory or inhibitory neuronal clusters.

#### Differential gene expression analysis

Differential gene expression analysis between WT an KO groups was performed using the Seurat function FindMarkers (min.pct = .00001, logfc.threshold = 0) with a Wilcoxon Rank Sum test. Genes with an adjusted p-value <.05 were considered differentially expressed between WT and KO.

#### Gene ontology

PANTHER^67,68^ (v18.0) was used to perform an overrepresentation test against the biological process complete ontology using default parameters. All genes detected by FindMarkers was used as a background gene list. For conciseness and visualization, parent terms were excluded and only the most specific GO terms were plotted.

### Electrophysiology

Mice were deeply anesthetized and trans-cardially perfused with ice-cold aCSF containing (in mM): 124 NaCl, 2.5 KCl, 1.2 HaH2PO4, 24 NaHCO3, 5 HEPES, 13 Glucose, 1.3 MgSO4, 2.5 CaCl2. After perfusion, the brain was quickly removed, submerged and coronally sectioned on a vibratome (VT1200s, Leica) at 400 μm thickness in ice-cold aCSF. Slices were transferred to NMDG based recovery solution at 32°C of the following composition (in mM): 92 NMDG, 2.5 KCl, 1.2 NaH2PO4, 30 NaHCO3, 20 HEPES, 25 Glucose, 5 Sodium ascorbate, 2 Thiourea, 3 Sodium pyruvate, 10 MgSO4, 0.5 CaCl2. After 12-15 minutes recovery, slices were transferred to room temperature aCSF chamber (20-22°C) and left for at least 1 hour before recording. Following recovery, slices were placed in a recording chamber, fully submerged at a flow rate of 1.4∼1.6 mL/min and maintained at 29-30°C in oxygenated (95% O2, 5% CO2) aCSF, with 100μM Picrotoxin included.

In extracellular recordings, recording pipettes were fabricated by pulling borosilicate glass (World Precision Instruments, TW150-3). These pipettes exhibited a tip resistance ranging from 4 to 5 MΩ when filled with an aCSF. Stimulus pipettes were made by pulling theta glasses (TG150-4, Warner Instrument) and adjusting the tip size to 30∼50 um. For measurements of evoked post-synaptic response in Schaffer-collateral pathway, Hippocampal layers were identified using IR-DIC optics (BX51, Olympus), with visual guidance. The stimulus electrode was positioned on the surface of the striatum radiatum (s.r.) from the CA3 direction. A recording electrode was placed on the postsynaptic site in CA1, opposite the stimulus electrode.

Brief electrical pulses (0.2ms) were delivered on each sweep to measure post-synaptic responses every 10 seconds. Once responses stabilized, the intensity of the stimulus was sequentially adjusted for input-output (I-O) measurement. Voltage (V) ranges were from 3V to 30V in increments of 3V. We identified the field excitatory post-synaptic potential (fEPSP) based on the temporal separation of the fiber volley, and then averaged measured the fEPSP slope of the 10-90% range from baseline to peak. Ten responses at the same input were averaged.

Recordings were performed using a MultiClamp 700B (Molecular Devices) and Igor7 (WaveMetrics; recording artist addon, developed by Richard C Gerkin, github : https://github.com/rgerkin/recording-artist), filtered at 2.8 kHz and digitized at 10 kHz. Axon terminals were stimulated with brief (0.2 ms) pulses using isoflex isolator (Voltage control). Data were analyzed using Igor7.

### Behavioral assays

#### Behavioral cohorts

Male and female WT and KO mice were tested in the behavioral tests described below. Mice were 3-4 months old at the onset of behavioral testing. One cohort of mice was used for olfaction, social choice, and fear conditioning testing, in that order. A separate cohort of mice was used for NOR and T-maze testing. During testing, the experimenter was blinded to genotype of the mice.

#### Olfactory habituation/dishabituation test

Mice were tested for olfactory habituation & dishabituation using a published protocol^34^. In brief, mice received sequential presentations of cotton swabs scented with different odors in the following sequence: water, almond, vanilla, same sex conspecific, opposite sex conspecific. Each odor was presented in three consecutive trials for 2 min, with an intertrial interval of 1 min. Time spent interacting with each scented swab was manually analyzed by three scorers who were blinded to sex and genotype.

#### NOR/open field

Novel object recognition and open field were performed as described in Korb 2015^81^ and Korb 2017^82^. In brief, mice were placed in an empty arena for 10 minutes for a habituation period that also served as an open field assessment. One day later mice were habituated for an additional 2 min and briefly removed from the arena while two identical objects (either a faucet, a plastic pyramid, a small fish figurine, or stacked Legos) were placed in the box and mice were given 10 min to explore. On the following day, mice were returned to the box with one object they had previously seen and one new object in place of the original object and were allowed to explore for 10 min. All sessions were recorded using EthoVision^83^ software. Time spent interacting with each object was manually analyzed. Discrimination index was calculated as (time with novel object − time with familiar object)/(time with novel object + time with familiar object).

#### 3-chamber social choice assay

The social choice test was carried out in a three-chambered apparatus, consisting of a center chamber and two outer chambers. Before the start of the test and in a counter-balanced manner, one end chamber was designated the social chamber, into which a stimulus mouse would be introduced, and the other end chamber was designed the nonsocial chamber. Two identical, clear Plexiglas cylinders with multiple holes to allow for air exchange were placed in each end chamber. In the habituation phase of the test, the experimental mouse freely explores the three chambers with empty cue cylinders in place for 10 min. Immediately following habituation, an age- and sex-matched stimulus mouse was placed in the cylinder in the social chamber while a rock was simultaneously placed into the other cylinder in the nonsocial chamber. The experimental mouse was tracked during the 10 min habituation and 10 min social choice phases. All testing was recorded, and videos were analyzed using ANY-maze software.

#### T-maze

For T-maze testing mice were placed at the end of the center arm of a T shaped raised arena enclosed with plexiglass. Mice were allowed to freely explore for 7 minutes. Entries into each arm were defined as the full body of the mouse entering (not necessarily including the tail). A success was defined as a consecutive entry into each of the 3 arms without returning to the arm that the mouse had been in immediately prior. Successful triads over the total number of possible triads based on the total entries were calculated as correct triad/(total entries - 2).

#### Contextual and cued fear conditioning

Mice were handled for 2 minutes each on 3 consecutive days immediately prior to the onset of testing. On training day, mice were placed in individual chambers for 2 min followed by a loud tone lasting 30 s that co-terminated with a 2-s, 1.25-mA foot shock. One minute later mice received another tone-shock pairing and were then left undisturbed for an additional 1 min in the chamber before being returned to their home cage. Freezing behavior, defined as no movement except for respiration, was determined before and after the tone-shock pairings and scored by MedAssociates VideoFreeze software. To test for context-dependent learning, we placed mice back into the same testing boxes 24 hr later for a total of 5 min without any tone or shock, and again measured the total time spent freezing. Following an additional 24 hr, we tested for cue-dependent fear memory by placing the mice into a novel chamber consisting of altered flooring, wall-panel inserts, and vanilla scent. After 2 min in the chamber, the cue tone was played for a total of 3 min, and the total time spent freezing during the presentation of this cue tone was recorded. Long-term contextual and cued fear memory were again tested with the same protocol at 14 d (contextual) or 15d (cued) post-training.

## QUANTIFICATION AND STATISTICAL ANALYSIS

All statistical analyses were performed using readily available code in R or using GraphPad Prism. Number of replicates and details of statistical tests are reported in figure legends. Detailed information on statistical tests as well as all relevant test statistics can be found in Table S1. Details on statistical packages and parameters used can be found within each relevant methods section and the Key Resources table.

## SUPPLEMENTAL TABLE TITLES

**Supplemental Table 1. Detailed statistical information for all figures.**

**Supplemental Table 2. GO analysis of H2BE peaks related to Figure 1**.

**Supplemental Table 3. Complete DESeq2 results from RNA sequencing of H2BE WT and KO cortical neurons, related to Figure 4**.

**Supplemental Table 4. GO analysis of genes downregulated in H2BE KO cortical neurons, related to Figure 4.**

**Supplemental Table 5. GO analysis of genes upregulated in H2BE KO cortical neurons, related to Figure 4.**

**Supplemental Table 6. SynGO analysis of genes downregulated in H2BE KO cortical neurons, related to Figure 4**.

**Supplemental Table 7. SynGO analysis of genes upregulated in H2BE KO cortical neurons, related to Figure 4**.

**Supplemental Table 8. GO analysis of genes downregulated in H2BE-KO excitatory cortical neurons from cluster Ex_L2/3_2, related to Figure 5**.

**Supplemental Table 9. GO analysis of genes upregulated in H2BE-KO excitatory cortical neurons from cluster Ex_L2/3_2, related to Figure 5**.

**Supplemental Table 10. GO analysis of genes downregulated in H2BE-KO inhibitory cortical neurons from cluster Inh_Phactr1, related to Figure 5**.

**Supplemental Table 11. GO analysis of genes upregulated in H2BE-KO inhibitory cortical neurons from cluster Inh_Phactr1, related to Figure 5**.

