## Supplemental Figures 1-6 and Supplemental Table 1 for "Histone variant H2BE enhances chromatin accessibility in neurons to promote synaptic gene expression and long-term memory"

### Supplemental Figure 1

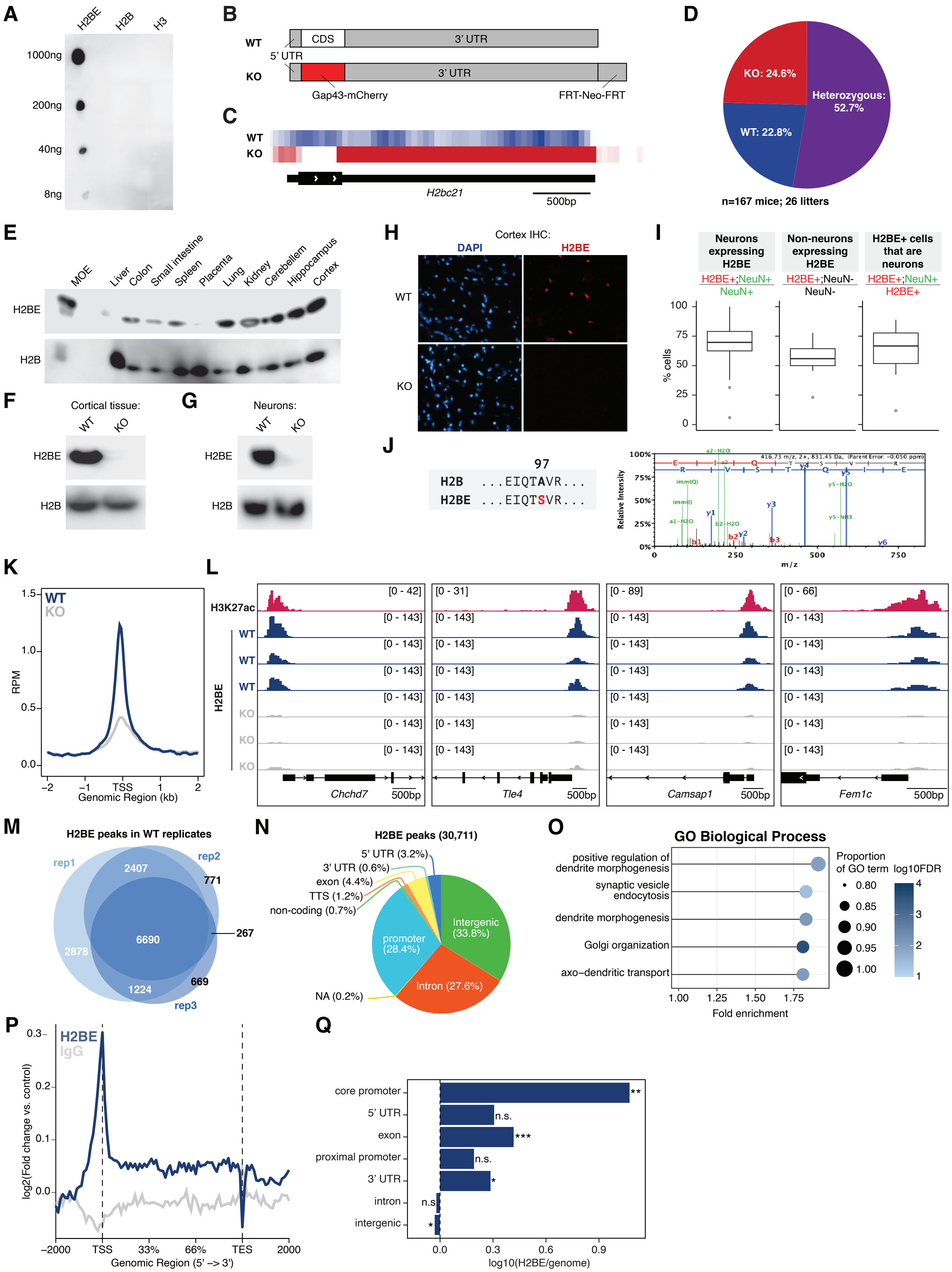

**Supplemental Figure 1.** (A) Dot blot analysis of H2BE antibody against N-terminal peptides for H2BE, H2B, and H3. (B) Schematic of the H2BE-KO/Gap43-mCherry KI allele as generated by Santoro & Dulac (2012). (C) RNA-sequencing heat map of WT and KO primary cortical neurons at the H2bc21 locus shows no reads in the H2BE coding sequence in KO neurons. (D) Pie chart of progeny from H2BE+/- x H2BE+/- crosses show expected Mendelian ratios (n = 126 mice from 26 litters). (E) Immunoblot for H2BE and H2B in various tissues from WT and KO mice. (F) Immunoblot for H2BE and H2B in cortical tissue. WT and KO demonstrates anti-H2BE antibody specificity. H2B serves as loading control. (G) Immunoblot for H2BE and H2B in primary cortical neurons. WT and KO demonstrates anti-H2BE antibody specificity. H2B serves as loading control. (H) Immunofluorescent images of cortical tissue from mice stained with H2BE and the nuclear marker DAPI demonstrates anti-H2BE antibody specificity. Scale bar = 30$\mu$m. (I) Quantification of immunofluorescent images shown in Figure 1C. (J) Representative MS/MS spectra of the H2B and H2BE peptides. (K) Metaplot comparison of CUT&Tag sequencing average signal from WT and KO primary cortical neurons demonstrates anti-H2BE antibody specificity. Plot shows read counts per million mapped reads (RPM) around the transcription start site (TSS) +/- 2kb (n = 1 biological replicate per genotype). (L) H2BE CUT&Tag genome browser tracks at select gene promoters with high enrichment of H2BE in WT. Loss of signal in KO neurons demonstrates anti-H2BE antibody specificity. H3K27ac CUT&Tag genome browser tracks were included to define nucleosome positions. (M) Overlap analysis of H2BE peaks detected by CUT&Tag from 3 biological replicates. (N) Genomic distribution of H2BE peaks detected by CUT&Tag. (M) Gene ontology enrichment analysis of genes where H2BE is enriched. (P) Metaplot of H2BE and control IgG ChIP-seq in WT primary cortical neurons. Plot shows read counts per million mapped reads (RPM) between the transcription start site (TSS) and transcription end site (TES) +/- 2kb (n = 2 biological replicates per genotype). (Q) Genomic distribution of ChIP-seq H2BE enrichment sites relative to the mouse genome (Chi-square test). *p<.05,**p<.01,***p<.001

#### Supplemental Figure 2

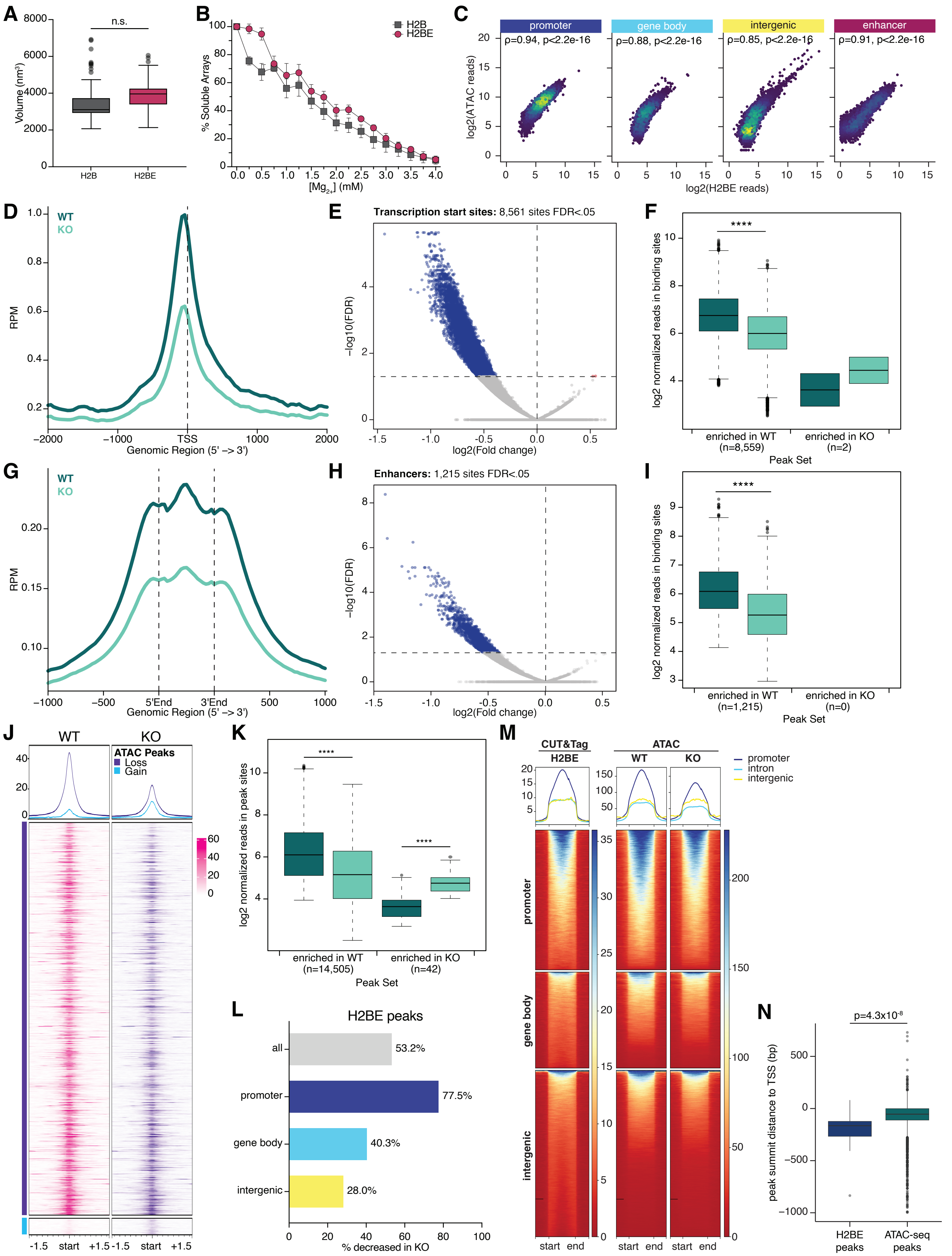

**Supplemental Figure 2.** (A) Volume of H2B or H2BE nucleosome arrays. (n = 60 H2B arrays, 40 H2BE arrays; Welch's t-test). Volume = total pixel intensity of each array. (B) $Mg^{2+}$ precipitation assay for chromatin arrays containing canonical H2B or variant H2BE (n = 4 per group). (C) Correlation between H2BE reads in WT neurons and changes in ATAC-seq signal between WT and KO at promoters, gene body, intergenic regions, and enhancers. (D) Metaplot comparison of ATAC-seq average signal from WT and KO primary cortical neurons at all transcription start sites. Plot shows read counts per million mapped reads (RPM) around the transcription start site (TSS) +/- 2kb (n = 3 WT biological replicates, 4 KO biological replicates). (E) Volcano plot showing differential chromatin accessibility between WT and KO at transcription start sites. Blue = peaks that decrease in KO; red = peaks that increase in KO; gray = peaks below significance cut-off of FDR<.05. (F) Normalized ATAC-seq read counts at transcription start sites with significantly different accessibility in WT (left) or KO (right) (unpaired t-test). (G) Metaplot comparison of ATAC-seq average signal from WT and KO primary cortical neurons at all enhancers identified in brain tissue. Plot shows read counts per million mapped reads (RPM) at all measured enhancers +/- 1kb (n = 3 WT biological replicates, 4 KO biological replicates). (H) Volcano plot showing differential chromatin accessibility between WT and KO at enhancers. Blue = peaks that decrease in KO; red = peaks that increase in KO; gray = peaks below significance cut-off of FDR<.05. (I) Normalized ATAC-seq read counts at transcription start sites with significantly different accessibility in WT (left) or KO (right). (unpaired t-test). (J) Summary plot (top) of peaks with decreased (purple) or increased (blue) accessibility in KO with heatmap (bottom) showing each TSS. The majority of TSS regions show decreased accessibility in KO. (K) Normalized ATAC-seq read counts at all WT ATAC-seq peaks with significantly different accessibility in WT (left) or KO (right). (unpaired t-test). (L) Percent of all detected ATAC-seq peaks within H2BE binding sites at promoters, within the gene body, or intergenic that are significantly decreased in KO. (M) Summary plot (top) and heatmap (bottom) of CUT&Tag reads and ATAC-seq reads at all H2BE binding sites, partitioned by genomic region. (N) Distance of ATAC-seq peak summits and H2BE peak summits to the TSS of that gene show that H2BE peaks are further upstream from the TSS (unpaired t-test). ****p<.0001

#### Supplemental Figure 3

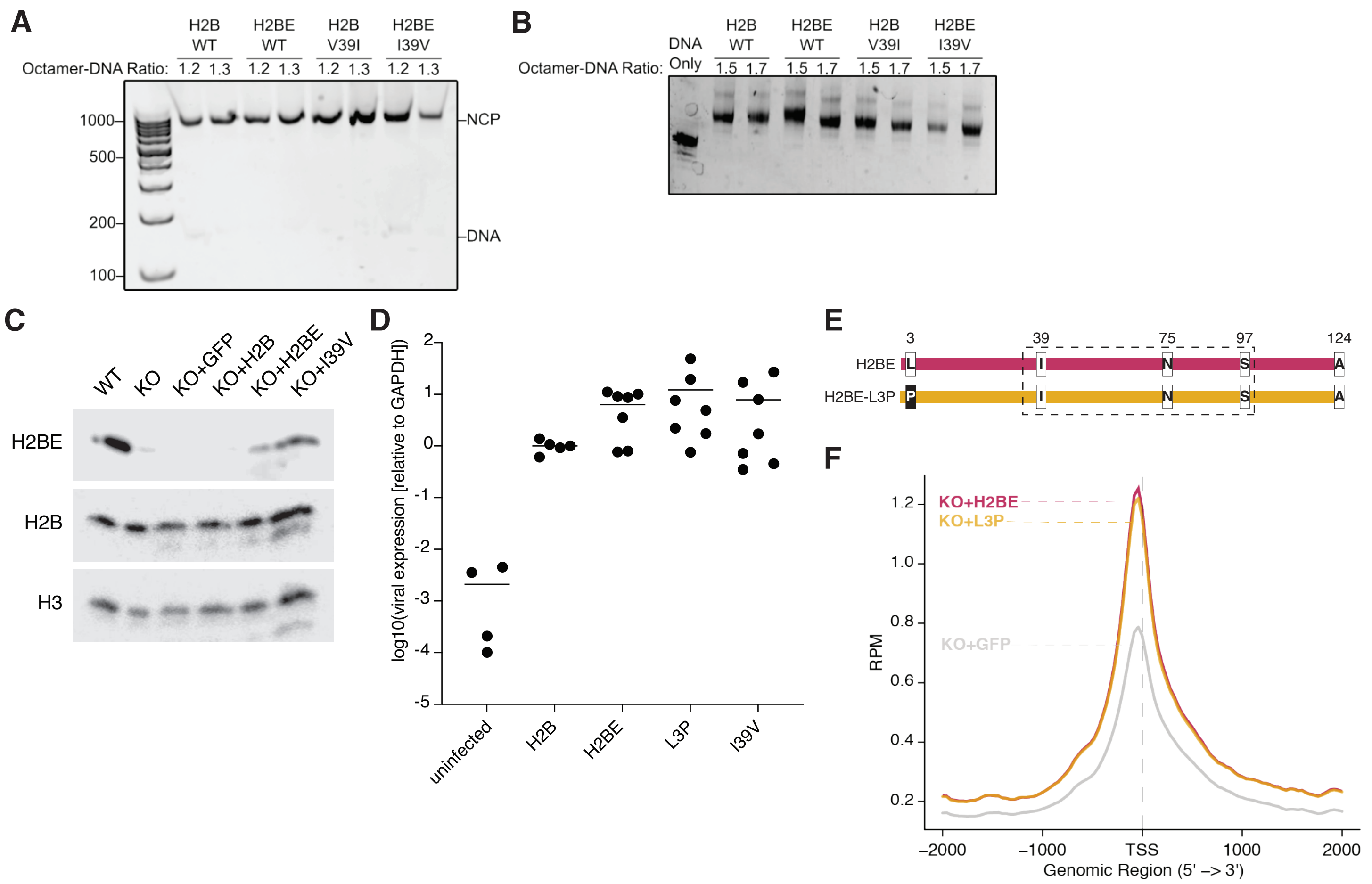


**Supplemental Figure 3.** (A) Mononucleosome electrophoretic mobility shift assay using native polyacrylamide gel electrophoresis to validate nucleosome assembly, stained using ethidium bromide to visualize DNA. (B) Chromatin array electrophoretic mobility shift assay using native agarose-polyacrylamide gel electrophoresis (APAGE) to validate chromatin array assembly, stained with ethidium bromide. (C) Western blot for H2BE and H2B in protein lysates from WT and KO neurons, and KO neurons infected with viruses expressing WT H2BE, H2BE-I39V, canonical H2B, or a GFP control. Statistical details in Table S1. (D) Relative expression of viral genes in primary neurons infected with viruses expressing WT H2BE, canonical H2B, or H2BE-L3P quantified by qRT-PCR. Expression is normalized to GAPDH. (E) Schematic WT H2BE and mutant H2BE-L3P sequences. (F) ATAC-seq average signal at all genes (n = 3-8 biological replicates per condition). Plot shows read counts per million mapped reads (RPM) around the transcription start site (TSS) +/- 2kb.

### Supplemental Figure 4

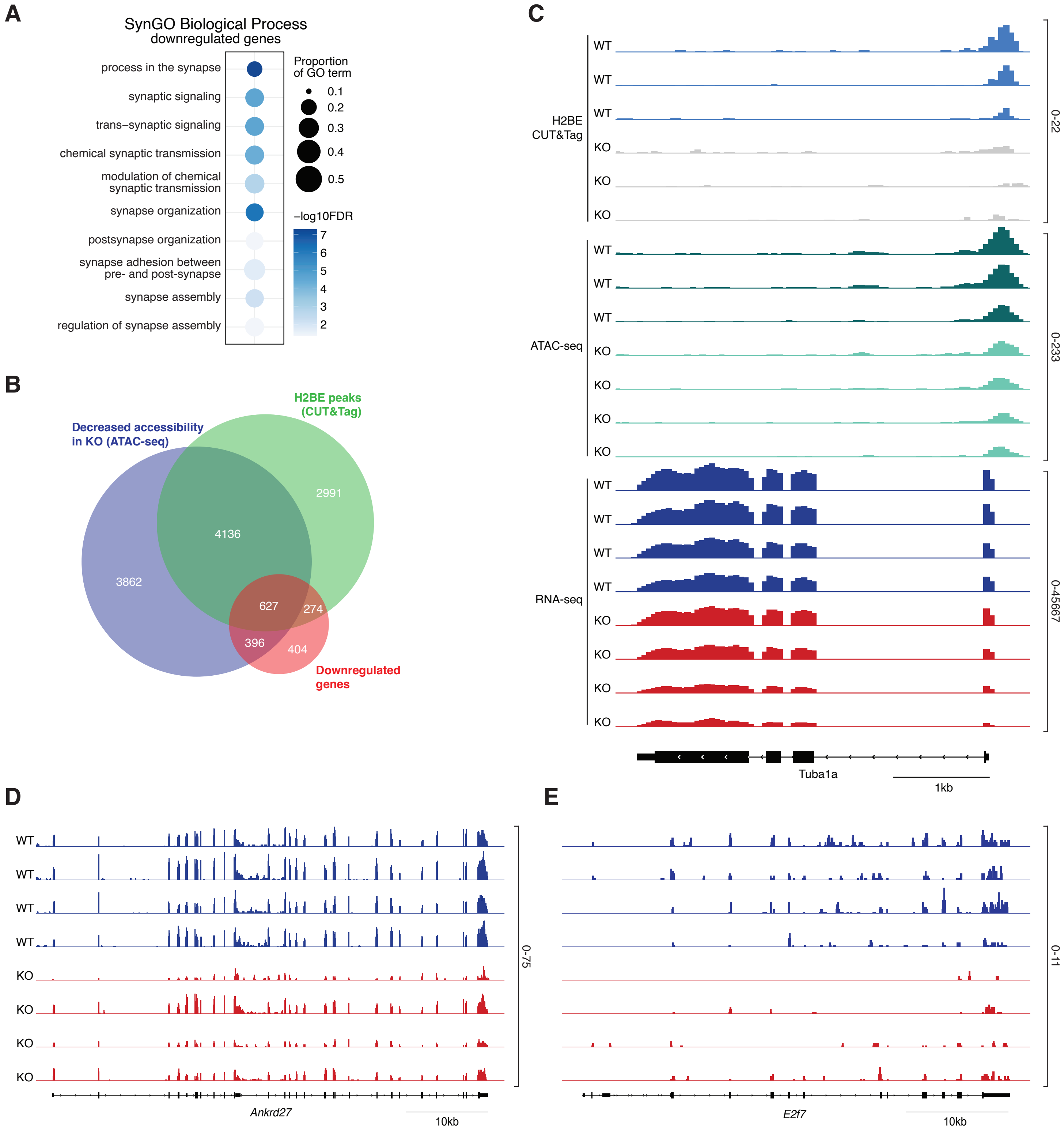


**Supplemental Figure 4.** (A) Synaptic gene ontology enrichment analysis by SynGO of genes where H2BE is enriched. (B) Overlap analysis of genes with H2BE enrichment, decreased accessibility in KO, and decreased transcription in KO. (C) CUT&Tag, ATAC-seq, and RNA-seq genome browser tracks at the Tuba1a locus in WT and KO cortical neurons. (D-E) RNA-seq tracks from WT and KO primary cortical neurons at (D) Ankrd27 and (E) E2f7 previously shown in Figs 1 and 2.

#### Supplemental Figure 5

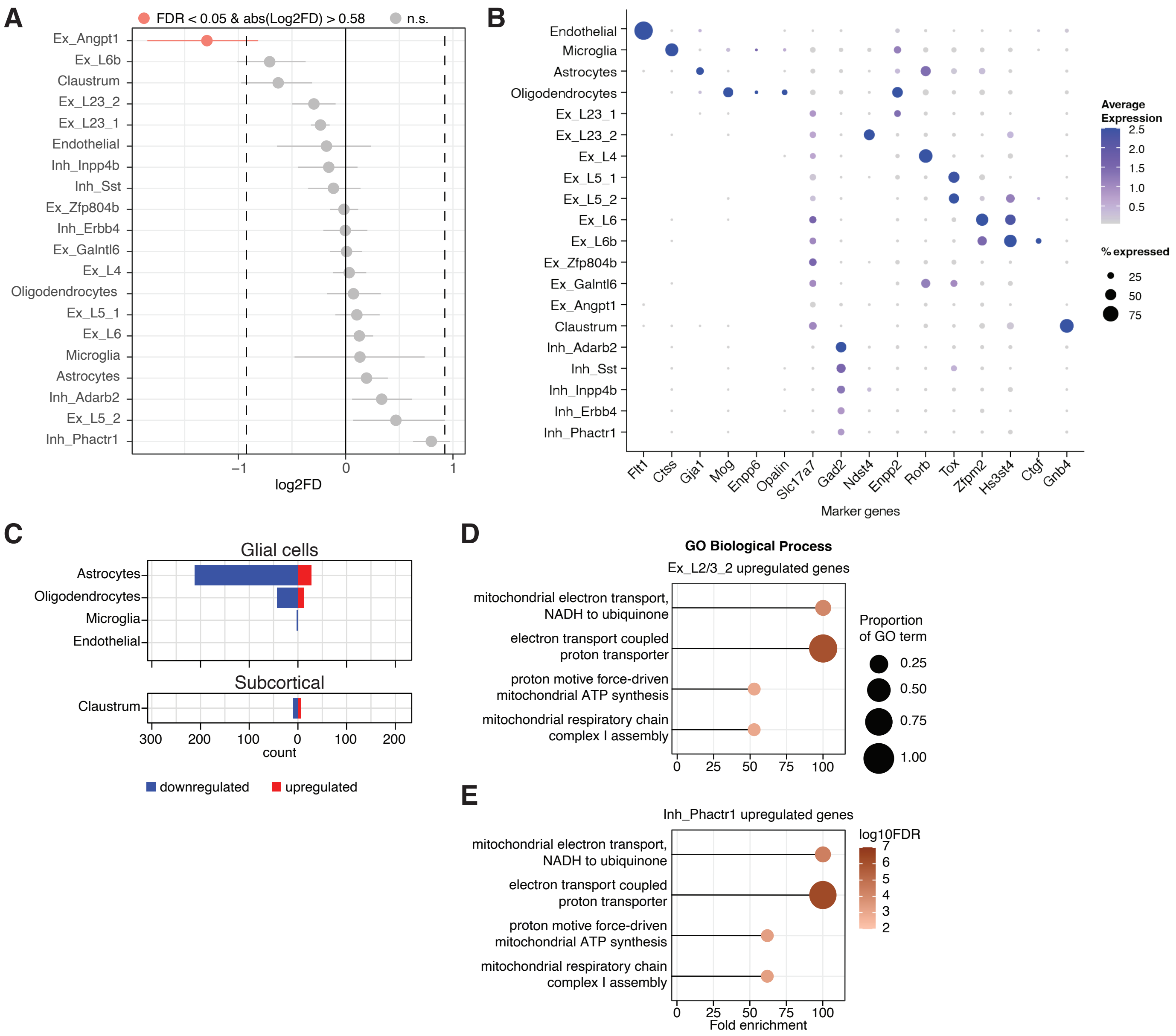


**Supplemental Figure 5.** (A) Analysis of the proportion of WT and KO nuclei in each cluster. Data represents the fold change in WT vs KO nuclei in each cluster and a confidence interval for the magnitude difference. Clusters with fold difference >1.8 and FDR<.05 highlighted in pink. Ex_Angpt1 cluster contains only small number of nuclei with 84 nuclei in WT and 45 in KO. (B) Bubble plot showing expression of cell type-specific marker genes for each cortical cell cluster. (C) Number of differentially expressed genes within glial clusters and claustral neurons from single-nucleus RNA-seq. (D-E) CUT&Tag, ATAC-seq, and RNA-seq genome browser tracks at the Tuba1a locus in WT and KO cortical neurons. Note that these gene ontology analyses produced the same results because of a high concordance between neuronal clusters and by nature of the small gene lists produced by snRNA-seq.

### Supplemental Figure 5

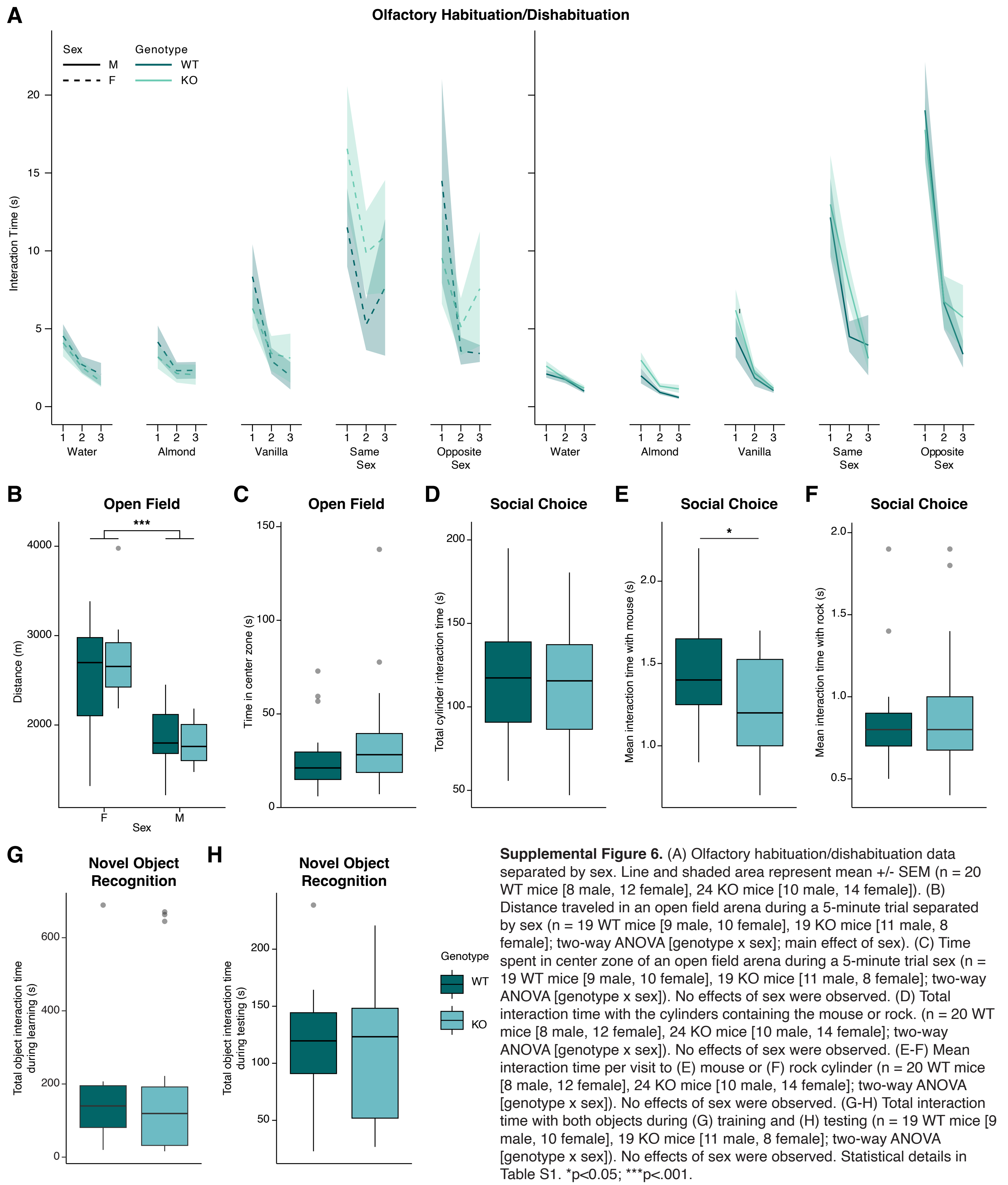


**Supplemental Figure 6.** (A) Olfactory habituation/dishabituation data separated by sex. Line and shaded area represent mean +/- SEM (n = 20 WT mice [8 male, 12 female], 24 KO mice [10 male, 14 female]). (B) Distance traveled in an open field arena during a 5-minute trial separated by sex (n = 19 WT mice [9 male, 10 female], 19 KO mice [11 male, 8 female]; two-way ANOVA [genotype x sex]; main effect of sex). (C) Time spent in center zone of an open field arena during a 5-minute trial sex (n = 19 WT mice [9 male, 10 female], 19 KO mice [11 male, 8 female]; two-way ANOVA [genotype x sex]). No effects of sex were observed. (D) Total interaction time with the cylinders containing the mouse or rock. (n = 20 WT mice [8 male, 12 female], 24 KO mice [10 male, 14 female]; two-way ANOVA [genotype x sex]). No effects of sex were observed. (E-F) Mean interaction time per visit to (E) mouse or (F) rock cylinder (n = 20 WT mice [8 male, 12 female], 24 KO mice [10 male, 14 female]; two-way ANOVA [genotype x sex]). No effects of sex were observed. (G-H) Total interaction time with both objects during (G) training and (H) testing (n = 19 WT mice [9 male, 10 female], 19 KO mice [11 male, 8 female]; two-way ANOVA [genotype x sex]). No effects of sex were observed. Statistical details in Table S1. *p<0.05; ***p<.001.

**Supplemental Table 1. Detailed statistical information for all figures.**

| figure | exp_descrip | n | test | covariates | pval | test_stat1 | test_stat1_value | test stat 2 | test stat 2 value |
|---|---|---|---|---|---|---|---|---|---|
| 1D | mass spec | P0: 5M, 5F biological replicates | 2-way ANOVA | Interaction | 0.9921 | F | F (4, 28) = 0.06358 | | |
| | | 4w: 3M, 3F biological replicates | | Age | <0.0001 | F | F (4, 28) = 20.20 | | |
| | | 8w: 3M, 3F biological replicates | | Sex | 0.989 | F | F (1, 28) = 0.0001950 | | |
| | | 12w: 5M, 5F biological replicates | Šídák's multiple comparisons test | P0 vs. 4w | 0.5276 | t | 1.405 | DF | 28 |
| | | 16w: 5M, 5F biological replicates | | P0 vs. 8w | 0.0027 | t | 3.822 | DF | 28 |
| | | | | P0 vs. 12w | 0.0011 | t | 4.172 | DF | 28 |
| | | | | P0 vs. 16w | <0.0001 | t | 8.529 | DF | 28 |
| 1F | CUT&Tag genomic distribution | 3 WT biological replicates | Chi-square test of independence for each partition | core promoter | <.001 | | | | |
| | | 3 KO biological replicates | (categorizing regions as observed/expected) | 5' UTR | <.001 | | | | |
| | | | | exon | <.001 | | | | |
| | | | | proximal promoter | <.001 | | | | |
| | | | | 3' UTR | <.001 | | | | |
| | | | | intron | 0.01<=p<0.05 | | | | |
| | | | | intergenic | <.001 | | | | |
| 1I | CUT&Tag peak score by expression | 3 WT biological replicates | one-way ANOVA | expression | <2E-16 | F | F(4) = 1006 | | |
| | | 3 KO biological replicates | pairwise t-tests with Bonferroni correction | high vs. mid | 1.40E-14 | | | | |
| | | | | high vs. low | <2E-16 | | | | |
| | | | | high vs. non | <2E-16 | | | | |
| | | | | high vs. ko_all | <2E-16 | | | | |
| | | | | mid vs. low | <2E-16 | | | | |
| | | | | mid vs. non | <2E-16 | | | | |
| | | | | mid vs. ko_all | <2E-16 | | | | |
| | | | | low vs. non | 0.00023 | | | | |
| | | | | low vs. ko_all | <2E-16 | | | | |
| | | | | non vs. ko_all | 0.72167 | | | | |
| 2A | thermal stability | 3 H2B nucleosome preps | Welch's unpaired t-tests with Holm-Šídák method | Tm1 | 0.889132 | t ratio | 0.1425 | df | 11.7 |
| | | 3 H2BE nucleosome preps | | | | | | | |
| | | 3 technical replicates per nuc | | Tm2 | 0.141806 | t ratio | 1.945 | df | 13.09 |
| 2C | AFM V/SA | 60 H2B arrays | Two-tailed unpaired Welch's t-test | | | | | | |
| | | 40 H2BE arrays | | variant | <0.0001 | t | 8.606 | df | 74.88 |
| 2F | ATAC reads - WT ATAC peaks | 3 WT biological replicates | unpaired t-test | | | | | | |
| | | 4 KO biological replicates | | genotype | <0.0001 | | | | |
| 2J | ATAC reads - H2BE peaks | 3 WT biological replicates | unpaired t-test | | | | | | |
| | | 4 KO biological replicates | | genotype | <0.0001 | | | | |
| 2L | ATACvH2BE | ATAC: | Spearman correlation | | | | | | |
| | | 3 WT biological replicates | | | | | | | |
| | | 4 KO biological replicates | | | | | | | |
| | | CUT&Tag: | | | | | | | |
| | | 3 WT biological replicates | | | | | | | |
| | | 3 KO biological replicates | | | <2.2E-16 | ρ | 0.742955 | | |
| 3C | thermal stability | 3 H2B nucleosome preps | two-way ANOVA | Tm | <0.0001 | F | F (1, 19) = 6625 | df | 11.7 |
| | | 3 H2BE nucleosome preps | | | | | | | |
| | | 3 H2B-V39I nucleosome preps | | | | | | | |
| | | 3 H2BE-I39V nucleosome preps | | | | | | | |
| | | 3 technical replicates per nucleosome | | Variant | 0.0865 | F | F (3, 19) = 2.546 | df | 13.09 |
| 4C | RNA read counts by deseq list | 4 WT biological replicates | one-way ANOVA | list (down, up, non-deg) | 1.84E-09 | F | F(2) = 20.14 | | |
| | | 4 KO biological replicates | pairwise t-tests with Bonferroni correction | down vs. up | 1.10E-07 | | | | |
| | | | | down vs. non-deg | 5.80E-09 | | | | |
| | | | | up vs. non-deg | 0.48 | | | | |
| 4E | ATAC log2FC by deseq list | 3 WT biological replicates | one-way ANOVA | list (down, up, non-deg) | 2.46E-11 | F | F(2) = 24.46 | | |
| | | 4 KO biological replicates | pairwise t-tests with Bonferroni correction | down vs. up | 4.20E-08 | | | | |
| | | | | down vs. non-deg | 3.60E-11 | | | | |
| | | | | up vs. non-deg | 1 | | | | |
| 6A | fEPSP | 5 WT mice (2M, 3F) | simple linear regression | WT | | r2 | 0.6748 | | |
| | | 4 KO mice (2M, 2F) | | KO | | r2 | 0.5006 | | |
| | | 11 WT slices | repeated measures two-way ANOVA | Interaction | 3.8E-11 | F | F (8, 176) = 9.733 | | |
| | | 13 KO slices | | Stim | 1E-12 | F | F (8, 176) = 189.9 | | |
| | | | | Genotype | 0.009401589 | F | F (1, 22) = 8.099 | | |

| | | | | | | | | | |
|---|---|---|---|---|---|---|---|---|---|
| | | | | Subject | 1E-12 | F | F (22, 176) = 36.76 | | |
| | | | Šídák's multiple comparisons test | 3 | 0.999999911 | t | 0.2082 | df | 198 |
| | | | | 6 | 0.977292717 | t | 0.9499 | df | 198 |
| | | | | 9 | 0.587360621 | t | 1.684 | df | 198 |
| | | | | 12 | 0.204294415 | t | 2.257 | df | 198 |
| | | | | 15 | 0.027224398 | t | 2.998 | df | 198 |
| | | | | 18 | 0.011268778 | t | 3.272 | df | 198 |
| | | | | 21 | 0.003448716 | t | 3.613 | df | 198 |
| | | | | 24 | 0.000916007 | t | 3.967 | df | 198 |
| | | | | 27 | 0.000283598 | t | 4.261 | df | 198 |
| 6B | fiber volley | 5 WT mice (2M, 3F) | simple linear regression | WT | | r2 | 0.7431 | | |
| | | 4 KO mice (2M, 2F) | | KO | | r2 | 0.7241 | | |
| | | 11 WT slices | repeated measures two-way ANOVA | Interaction | 0.132775292 | F | F (9, 198) = 1.550 | | |
| | | 13 KO slices | | Stim | <0.000000000001 | F | F (9, 198) = 149.8 | | |
| | | | | Genotype | 0.840472436 | F | F (1, 22) = 0.04149 | | |
| | | | | Subject | <0.000000000001 | F | F (22, 198) = 11.72 | | |
| | | | Šídák's multiple comparisons test | 3 | >0.999999999999 | t | 0.02492 | df | 220 |
| | | | | 6 | 0.999999915 | t | 0.2488 | df | 220 |
| | | | | 9 | 0.999967261 | t | 0.4628 | df | 220 |
| | | | | 12 | 0.99979824 | t | 0.5646 | df | 220 |
| | | | | 15 | 0.999460935 | t | 0.6308 | df | 220 |
| | | | | 18 | 0.999714318 | t | 0.587 | df | 220 |
| | | | | 21 | 0.999881457 | t | 0.5323 | df | 220 |
| | | | | 24 | 0.999933316 | t | 0.4998 | df | 220 |
| | | | | 27 | 0.999999965 | t | 0.2272 | df | 220 |
| 6D | olfaction | 20 WT mice (8M, 12 F) | three-way ANOVA (water) | Trial | 1.02E-08 | F | 37.146 | df | 1 |
| | | 24 KO mice (10M, 14F) | | Genotype | 0.9975 | F | 0 | df | 1 |
| | | | | Sex | 4.66E-06 | F | 22.724 | df | 1 |
| | | | | Trial:Genotype | 0.7341 | F | 0.116 | df | 1 |
| | | | | Trial:Sex | 0.0427 | F | 4.183 | df | 1 |
| | | | | Genotype:Sex | 0.2119 | F | 1.573 | df | 1 |
| | | | | Trial:Genotype:Sex | 0.8196 | F | 0.052 | df | 1 |
| | | | three-way ANOVA (almond) | Trial | 1.38E-05 | F | 20.312 | df | 1 |
| | | | | Genotype | 0.505349 | F | 0.446 | df | 1 |
| | | | | Sex | 0.000109 | F | 15.874 | df | 1 |
| | | | | Trial:Genotype | 0.993601 | F | 0 | df | 1 |
| | | | | Trial:Sex | 0.783725 | F | 0.076 | df | 1 |
| | | | | Genotype:Sex | 0.053856 | F | 3.781 | df | 1 |
| | | | | Trial:Genotype:Sex | 0.428519 | F | 0.631 | df | 1 |
| | | | three-way ANOVA (vanilla) | Trial | 1.29E-08 | F | 36.636 | df | 1 |
| | | | | Genotype | 0.548 | F | 0.363 | df | 1 |
| | | | | Sex | 0.0175 | F | 5.783 | df | 1 |
| | | | | Trial:Genotype | 0.8236 | F | 0.05 | df | 1 |
| | | | | Trial:Sex | 0.8062 | F | 0.06 | df | 1 |
| | | | | Genotype:Sex | 0.4634 | F | 0.541 | df | 1 |
| | | | | Trial:Genotype:Sex | 0.1155 | F | 2.51 | df | 1 |
| | | | three-way ANOVA (same sex) | Trial | 7.05E-05 | F | 16.795 | df | 1 |
| | | | | Genotype | 0.1071 | F | 2.63 | df | 1 |
| | | | | Sex | 0.0433 | F | 4.16 | df | 1 |
| | | | | Trial:Genotype | 0.6329 | F | 0.229 | df | 1 |
| | | | | Trial:Sex | 0.242 | F | 1.381 | df | 1 |
| | | | | Genotype:Sex | 0.2858 | F | 1.148 | df | 1 |
| | | | | Trial:Genotype:Sex | 0.9904 | F | 0 | df | 1 |
| | | | three-way ANOVA (opposite sex) | Trial | 5.17E-08 | F | 33.183 | df | 1 |
| | | | | Genotype | 0.8154 | F | 0.055 | df | 1 |
| | | | | Sex | 0.0895 | F | 2.924 | df | 1 |
| | | | | Trial:Genotype | 0.1145 | F | 2.522 | df | 1 |
| | | | | Trial:Sex | 0.0436 | F | 4.147 | df | 1 |
| | | | | Genotype:Sex | 0.9599 | F | 0.003 | df | 1 |
| | | | | Trial:Genotype:Sex | 0.466 | F | 0.534 | df | 1 |
| 6E | open field, distance | 19 WT mice (9M, 10F) | two-way ANOVA | Genotype | 0.921 | F | 0.01 | df | 1 |
| | | 19 KO mice (11M, 8F) | | Sex | 8.57E-06 | F | 27.382 | df | 1 |

| | | | | | | | | | |
|---|---|---|---|---|---|---|---|---|---|
| | | | | Genotype:Sex | 0.403 | F | 0.716 | df | 1 |
| 6F | social choice | 20 WT mice (8M, 12 F) | two-way ANOVA | Genotype | 0.2775 | F | 1.2119 | df | 1 |
| | | 24 KO mice (10M, 14F) | | Sex | 0.108 | F | 2.7022 | df | 1 |
| | | | | Genotype:Sex | 0.3282 | F | 0.9798 | df | 1 |
| 6G | t-maze | 19 WT mice (9M, 10F) | two-way ANOVA | Genotype | 0.925 | F | 0.009 | df | 1 |
| | | 19 KO mice (11M, 8F) | | Sex | 0.293 | F | 1.147 | df | 1 |
| | | | | Genotype:Sex | 0.602 | F | 0.278 | df | 1 |
| 6H | NOR | 19 WT mice (9M, 10F) | two-way ANOVA | Genotype | 0.0331 | F | 5.205 | df | 1 |
| | | 19 KO mice (11M, 8F) | | Sex | 0.5877 | F | 0.303 | df | 1 |
| | | | | Genotype:Sex | 0.4115 | F | 0.702 | df | 1 |
| 6I | cued FC | 20 WT mice (8M, 12 F) | two-way ANOVA (acquisition, tone1) | Genotype | 0.903 | F | 0.015 | df | 1 |
| | | 24 KO mice (10M, 14F) | | Sex | 9.62E-01 | F | 0.002 | df | 1 |
| | | | | Genotype:Sex | 0.337 | F | 0.945 | df | 1 |
| | | | two-way ANOVA (48h recall) | Genotype | 0.1084 | F | 2.696 | df | 1 |
| | | | | Sex | 0.2928 | F | 1.136 | df | 1 |
| | | | | Genotype:Sex | 0.0844 | F | 3.131 | df | 1 |
| | | | two-way ANOVA (15d recall) | Genotype | 0.00128 | F | 12.003 | df | 1 |
| | | | | Sex | 0.4123 | F | 0.686 | df | 1 |
| | | | | Genotype:Sex | 0.84133 | F | 0.041 | df | 1 |
| 6J | contextual FC | 20 WT mice (8M, 12 F) | two-way ANOVA (acquisition, pre-shock) | Genotype | 0.0639 | F | 3.633 | df | 1 |
| | | 24 KO mice (10M, 14F) | | Sex | 0.9885 | F | 0 | df | 1 |
| | | | | Genotype:Sex | 0.5271 | F | 0.407 | df | 1 |
| | | | two-way ANOVA (24h recall) | Genotype | 0.137 | F | 2.298 | df | 1 |
| | | | | Sex | 0.96 | F | 0.003 | df | 1 |
| | | | | Genotype:Sex | 0.563 | F | 0.341 | df | 1 |
| | | | two-way ANOVA (15d recall) | Genotype | 0.00735 | F | 7.979 | df | 1 |
| | | | | Sex | 0.40991 | F | 0.694 | df | 1 |
| | | | | Genotype:Sex | 0.25249 | F | 1.348 | df | 1 |
| supp_1Q | ChIP-seq genomic distribution | 2 WT biological replicates | Chi-square test of independence for each partition | core promoter | 0.001<=p<0.01 | | | | |
| | | 2 KO biological replicates | (categorizing regions as observed/expected) | 5' UTR | n.s. | | | | |
| | | | | exon | <.001 | | | | |
| | | | | proximal promoter | n.s. | | | | |
| | | | | 3' UTR | 0.01<=p<0.05 | | | | |
| | | | | intron | n.s. | | | | |
| | | | | intergenic | 0.01<=p<0.05 | | | | |
| supp_2A | AFM volume | 60 H2B arrays<br>40 H2BE arrays | Two-tailed unpaired t-test | variant | 0.082 | t | 1.757 | df | 98 |
| supp_2N | ChIP-seq peak summit distance to TSS | 2 WT biological replicates per target (H2BE, IgG) | unpaired t-test | distance (bp) | 4.30E-08 | | | | |
| supp_5B | open field, distance, by sex | 19 WT mice (9M, 10F) | two-way ANOVA | Genotype | 0.921 | F | 0.01 | df | 1 |
| | | 19 KO mice (11M, 8F) | | Sex | 8.57E-06 | F | 27.382 | df | 1 |
| | | | | Genotype:Sex | 0.403 | F | 0.716 | df | 1 |
| supp_5C | open field, center | 19 WT mice (9M, 10F) | two-way ANOVA | Genotype | 0.214 | F | 1.6 | df | 1 |
| | | 19 KO mice (11M, 8F) | | Sex | 0.594 | F | 0.29 | df | 1 |
| | | | | Genotype:Sex | 0.998 | F | 0 | df | 1 |
| supp_5D | social choice, total cyl time | 20 WT mice (8M, 12 F) | two-way ANOVA | Genotype | 0.9923 | F | 0 | df | 1 |
| | | 24 KO mice (10M, 14F) | | Sex | 0.2778 | F | 1.211 | df | 1 |
| | | | | Genotype:Sex | 0.0691 | F | 3.489 | df | 1 |
| supp_5E | social choice, mean mouse visit | 20 WT mice (8M, 12 F) | two-way ANOVA | Genotype | 0.0147 | F | 6.507 | df | 1 |
| | | 24 KO mice (10M, 14F) | | Sex | 0.6711 | F | 0.183 | df | 1 |
| | | | | Genotype:Sex | 0.1012 | F | 0.3149 | df | 1 |
| supp_5F | social choice, mean rock visit | 20 WT mice (8M, 12 F) | two-way ANOVA | Genotype | 0.475 | F | 0.521 | df | 1 |
| | | 24 KO mice (10M, 14F) | | Sex | 0.142 | F | 2.244 | df | 1 |
| | | | | Genotype:Sex | 0.852 | F | 0.035 | df | 1 |
| supp_5G | NOR, habituation interaction time | 19 WT mice (9M, 10F) | two-way ANOVA | Genotype | 0.6945 | F | 0.157 | df | 1 |
| | | 19 KO mice (11M, 8F) | | Sex | 0.0297 | F | 5.195 | df | 1 |
| | | | | Genotype:Sex | 0.1526 | F | 2.151 | df | 1 |
| supp_5H | NOR, test interaction time | 19 WT mice (9M, 10F) | two-way ANOVA | Genotype | 0.655 | F | 0.205 | df | 1 |
| | | 19 KO mice (11M, 8F) | | Sex | 0.287 | F | 1.194 | df | 1 |
| | | | | Genotype:Sex | 0.671 | F | 0.186 | df | 1 |
